# In-depth sequence-function characterization reveals multiple paths to enhance phenylalanine ammonia-lyase (PAL) activity

**DOI:** 10.1101/2021.06.06.447205

**Authors:** Vikas D. Trivedi, Todd C. Chappell, Naveen B. Krishna, Anuj Shetty, Gladstone G. Sigamani, Karishma Mohan, Athreya Ramesh, Pravin Kumar R., Nikhil U. Nair

**Author notes:** Equal contribution (Vikas D. Trivedi, Todd C. Chappell).

## Abstract

Phenylalanine ammonia-lyases (PALs) deaminate L-phenylalanine to *trans*-cinnamic acid and ammonium and have widespread application in chemo-enzymatic synthesis, agriculture, and medicine. In particular, the PAL from *Anabaena variabilis* (*Trichormus variabilis*) has garnered significant attention as the active ingredient in Pegvaliase®, the only FDA-approved drug treating classical phenylketonuria (PKU). Although an extensive body of literature exists on structure, substrate-specificity, and catalytic mechanism, protein-wide sequence determinants of function remain unknown, which limits the ability to rationally engineer these enzymes. Previously, we developed a high-throughput screen (HTS) for PAL, and here, we leverage it to create a detailed sequence-function landscape of PAL by performing deep mutational scanning (DMS). Our method revealed 79 hotspots that affected a positive change in enzyme fitness, many of which have not been reported previously. Using fitness values and structure-function analysis, we picked a subset of residues for comprehensive single- and multi-site saturation mutagenesis to improve the catalytic activity of PAL and identified combinations of mutations that led to improvement in reaction kinetics in cell-free and cellular contexts. To understand the mechanistic role of the most beneficial mutations, we performed QM/MM and MD and observed that different mutants confer improved catalytic activity via different mechanisms, including stabilizing first transition and intermediate states and improving substrate diffusion into the active site, and decreased product inhibition. Thus, this work provides a comprehensive sequence-function relationship for PAL, identifies positions that improve PAL activity when mutated and assesses their mechanisms of action.

## INTRODUCTION

Phenylalanine ammonia-lyases (EC 4.3.1.24; PALs) non-oxidatively deaminate L-phenylalanine (Phe) to *trans*-cinnamic acid (tCA), releasing ammonium (NH ^+^), and are widely found associated with secondary metabolism in plants, bacteria, and fungi^1^. They are part of a family of enzymes that contain the rare, autocatalytically forming 4-methylideneimidazole-5-one (MIO) adduct, which enables deamination without an exogenous cofactor such as pyridoxal 5-phosphate (PLP) and/or co-substrate(s)^2^. Biocatalytic applications for natural product and fine chemical synthesis, as well as therapeutic potential have driven the discovery, expression, characterization, and engineering of PALs^3–8^. In particular, the recent success translating PAL into an enzyme replacement therapy for phenylketonuria (PKU) management and potential use as a cancer therapy have further increased interest in engineering this class of enzymes^9–12^. While there is extensive literature on the structure and catalytic mechanism of PALs, and a general understanding of how residues in the substrate-binding pocket contribute to specificity and turnover, led by semi-rational and homology-guided mutagenesis studies^13–15^, there is poor understanding of how more distal residues affect function.

Generally, directed evolution can identify mutational hotspots using large mutant libraries, independent of their proximity to the active site. But there have only been two such studies with PALs – one study resulted in modest improvement in activity^16^ whereas the other, conducted by us, identified only residues within the active site^17^. Because both studies involved characterization of a limited subset of mutants following directed evolution, conclusions could only be drawn regarding very specific mutations at a few positions. Deep mutational scanning (DMS), an emerging approach to assess sequence-function relationships^18^, can help identify functional hotspots^19^, and when coupled with directed evolution, can accelerate and broaden engineering campaigns^20,21^. Specifically, DMS can provide a comprehensive map of sequence–function relationships to explore the protein fitness landscapes^19^, discover new functionally relevant sites^22^, improve molecular energy functions, and identify beneficial combinations of mutations for protein engineering^23^. With the increasing successful application of DMS,^6,24–37^ we felt a systematic study exploring complete sequence-function relationships would be useful to better understand and engineer PALs with enhanced activity.

Previously, we developed a growth-coupled enrichment for rapid screening of high-activity variants of AvPAL* (the double mutant C503S-C565S PAL from *Anabaena variabilis*, currently used to formulate the PKU drug Pegvaliase®) in *E. coli*^17^. After a single round of directed evolution using this growth-coupled enrichment, we identified active site mutations G218S and M222L that possessed ~1.8-fold improvement in turnover frequency (k_cat_). Here, we vastly expand upon our prior work. First, we provide a detailed sequence-function landscape of AvPAL*, using DMS to identify 79 hotspot residues that improve activity. Next, we picked seven hotspots for comprehensive single- or multi-site saturation mutagenesis to study their interactions and further enhance the catalytic activity. We noted that the beneficial mutations were not well-represented in the natural sequence diversity of homologous PAL enzymes. We observed that few mutations showed positive fitness with increasing number of co-mutating residues. We also found that the best combination of mutation among 7 sites were a double (T102E-M222L) and triple (T012E-M222L-D306G) mutant that displayed a ~2.5-fold improvement in the k_cat_ (and >3-fold increase in catalytic efficiency). To understand the mechanistic role of key mutations in hyperactive variants, we performed modelling studies (Quantum Mechanical, QM/MM, and Molecular Dynamics, MD, including metadynamics) and concluded that there are multiple pathways to enhance PAL catalytic activity, including, i) decreased root mean square fluctuation (RMSF) of substrate in the active site, ii) greater proximity of the substrate to catalytic residues, iii) stabilization of the substrate in the near attack conformation, iv) stabilization of the transition and intermediate states, and v) facilitated diffusion of the substrate to the active site. Based on the unique experimental and computational insights, we also created a variant T102E-M222L-N453S that displayed lower product inhibition and ~6-fold higher activity in a whole-cell context. In summary, this study significantly advances basic and applied enzymology of PALs, a heretofore understudied class of enzymes with a wide array of applications.

## RESULTS & DISCUSSION

### Overview

The overall workflow of this work is summarized in **Fig. 1.** Starting with a randomly mutagenized library, we first performed deep mutational scanning (DMS) of AvPAL* using a growth-based high-throughput screen (HTS) to evaluate the fitness – or change in relative frequency over the three passages – of each mutation. Briefly, we deep sequenced plasmid libraries from the naïve and each of the three enrichment passages to identify mutations that occurred at each position, calculated the change in occurrence frequency for every mutation in each of the enriched populations relative to the naïve library (fitness), and mapped them onto the protein sequence and structure. Using fitness, structural insights, and domain knowledge, we classified certain positions as mutational hotspots from which we generated site-saturation mutagenesis libraries of each position alone, or in combination. We then enriched these libraries using our HTS, as before, and identified additional variants that with further enhanced activity. Next, we performed MD, including metadynamics, and QM/MM studies to characterize the catalytic mechanism of AvPAL*, and assess the functional impact that these mutations have on its catalytic activity. Finally, we used data from all investigations to devise hyperactive AvPAL* variants.

**Fig. 1:**
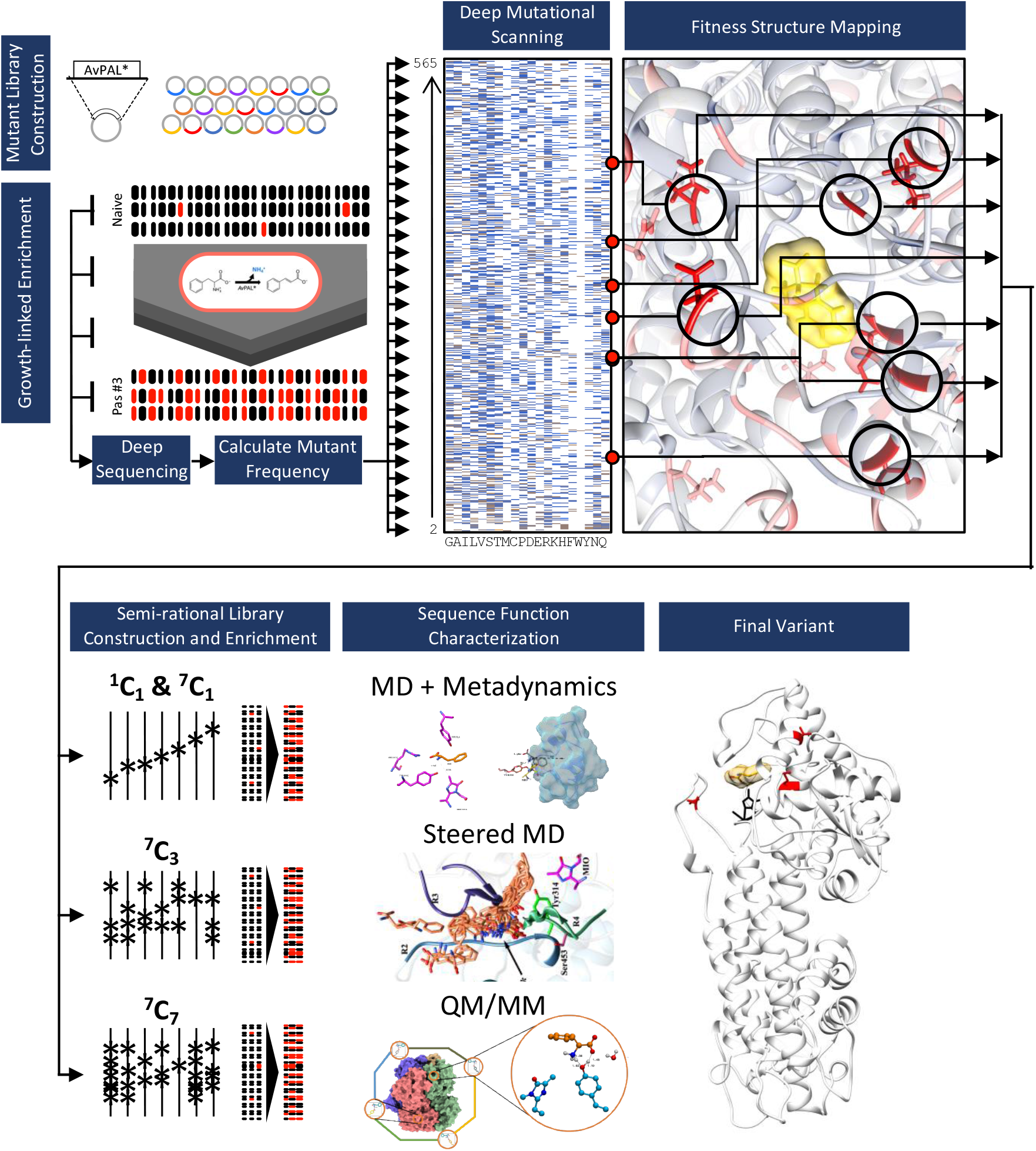
Overview of work. We started with a randomly mutagenized library of AvPAL* and performed deep mutational scanning (DMS). We identified mutational hotspots and used that to guide further mutagenesis and computational modeling studies (QM/MM, MD, and metadynamics) to assess multiple pathways to improve enzyme activity. Finally, culling data from DMS, targeted mutagenesis, and mechanistic modeling aided in design of hyperactive AvPAL* variants.

### Deep mutational scanning (DMS) of AvPAL* and analysis of active site residues

The naïve library contained approximately 2-4 amino acid mutations per gene, and had a broad distribution of mutations, with no major bias prior to enrichment (**Fig. 2a**). Since the naïve library was generated using error-prone PCR, it had an average of 5.6 substitutions per residue (not 20), with a range of 2−7 (**Fig. 2b**). Comparatively, the enriched library from the third passage averaged only 0.6 mutations per position, with a range of 0−5 mutations. The enriched library also contained only 222 positions with at least one mutation (relative to the overall protein size of 565 amino acids). We found that all premature nonsense codons in the naïve library were rapidly depleted, and the library also shifted from majority non-synonymous to synonymous mutations during enrichment (**Fig. S1**).

**Fig. 2:**
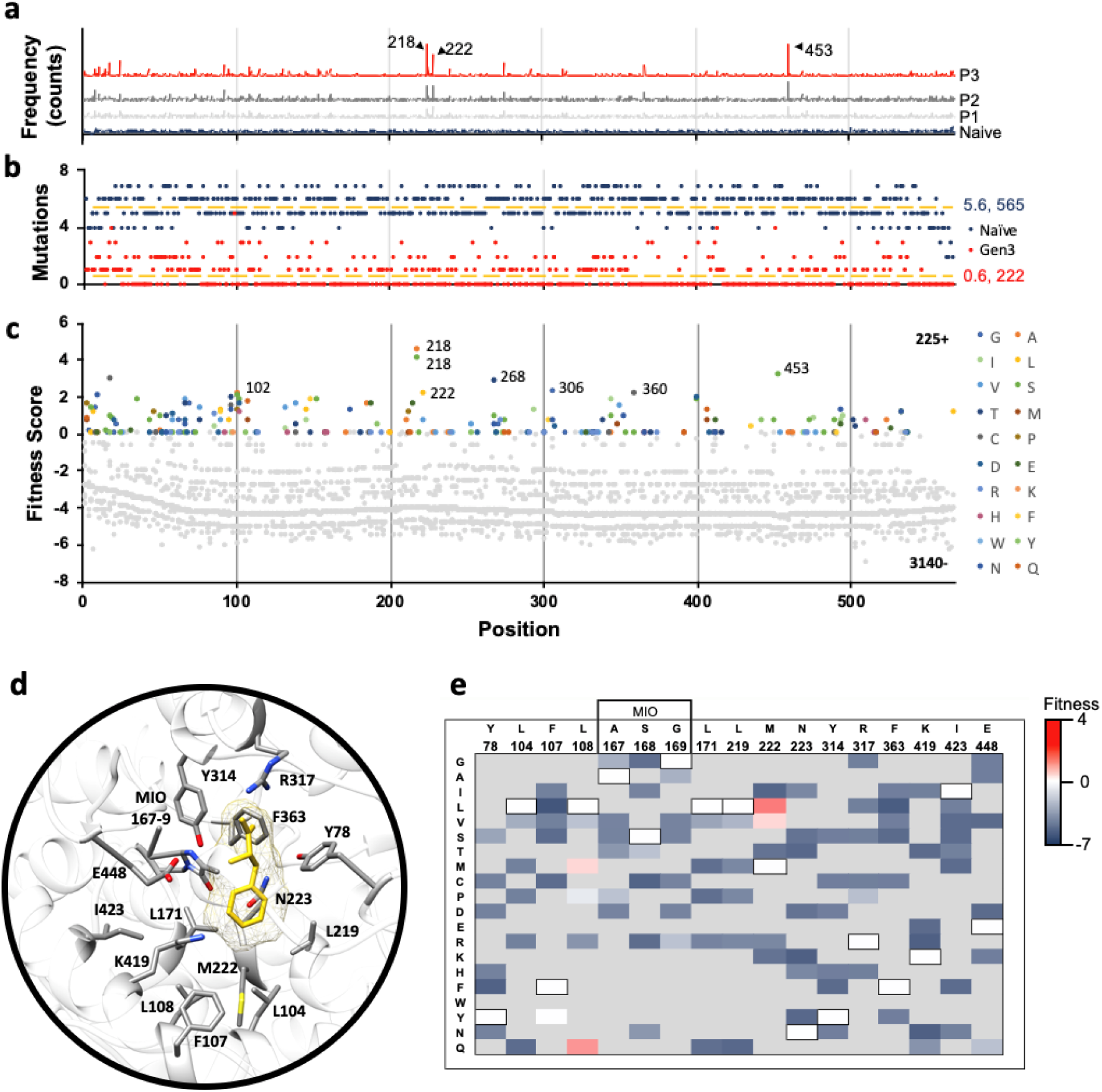
AvPAL* deep mutational scanning (DMS) outcomes. **a)** Mutation frequency in naïve and passages #1-3. Positions (a.a.) corresponding to the three most highly enriched positions are labeled (218, 222, and 453). **b)** Number of mutations sampled at each position in the naïve (dark blue) and passage #3 (red) libraries. The naïve library had an average of 5.6 mutations per position across all 565 positions. Passage #3 library had an average of 0.6 mutations per position across 222 positions. **c)** Fitness of all mutations present at all positions in the passage #3 library. Negative fitness denotes mutations that decrease in frequency over passages; the highest fitness mutations are labeled (G218A, G218S, M222L, I268T, D306G, G360C, and N453S). Position labels (x-axis) are the same for panels (a-c). **d)** Active site residues of AvPAL* (grey sticks) with phenylalanine ligand (yellow) docked. **e)** Fitness heatmap of active site residues at passage #3. Wildtype residues of AvPAL* are bordered in black and listed above the residue position. Grey boxes indicate mutations not sampled in library.

To evaluate the relative increase of each mutant variant in library during enrichment, we calculated a fitness score for each mutation. We found that the maximum fitness scores and growth rate across passages generally increased, as expected, indicative of strong positive selection for PAL activity (**Fig. S1–3**). We also found a good correlation between enzyme specific activity and fitness conferred (**Fig. S4**). Overall, 93% of mutations in the library had negative fitness in the third passage, indicating that most mutations are deleterious (**Fig. 2c**). Our data revealed that the active site residues – catalytic and substrate binding – were generally non-permissive to mutations (**Fig. 2d, e**). Only two positions had mutations with positive fitness (L108 and M222). Although we did not sample all 20 amino acids at each position, our data is in good agreement with published literature. Y78, Y314, and the MIO-forming triad (A167, S168, and G169) that are implicated to play essential roles in catalysis^38^ and highly conserved in PALs were found to be non-permissive to mutations. L104, F107, L108, and L171 in AvPAL* are part of the substrate binding pocket that interacts with the hydrophobic moiety of Phe. Among these, L104, L171, and L219 are three of the most highly conserved residues in tyrosine/phenylalanine ammonia lyases (TPALs) and mutases^2^ and any substitution had negative impact on fitness (**Fig. 2e**). F107 equivalent position shows more diversity, as PALs contains Phe^36^, TPALs contain basic amino acids Arg^37^ or His^30^, whereas ammonia mutase carry more polar amino acids Cys^28,39^ or Ser^40^. In PALs, F107 forms edge-edge interaction with the phenyl ring of the substrate^8^ and hence a mutation to Tyr is likely to be minimally disruptive, consistent with neutral fitness of F107Y in our data. In an earlier study, L108A, G mutations were shown to drastically reduce the enzyme activity suggesting it is not permissive to mutations^8^. However, we found L108Q, M to have positive fitness, suggesting the need for a large uncharged sidechain to fill-in the active site and maintain favorable interaction with the substrate. We confirmed L108Q and M to be more active and L108G to be less active than parental AvPAL* on phenylalanine (**Fig. S5**). Though L108M agrees with the requirement of hydrophobic residue to maintain nonpolar contacts^8^, L108Q is an unusual amino acid change to polar residue at otherwise conserved position. Generally, Leu at 108 equivalent position is conserved in ammonia-lyases active on phenylalanine, and His is found to be present in enzymes active on Tyr^2^. On performing sequence analysis on AvPAL* homologs against the RefSeq protein database, we observed out of the 998 sequences, Leu and His are present at the 108-equivalent position in 490 sequences each, accounting for 98.2% of the sequences, and Ala, Lys, Met, Gln, Thr are found in remaining 18 sequences (**Table S1**). The M222 position shows greater permissivity in the library and its natural diversity in homologs. Of the same 998 homologs, 469 had Val, 396 had Met, whereas, Ile, Asn, Thr, and Leu were found in 105, 15, 7 and 6 sequences, respectively. We found M222L, V to have higher fitness and thus, higher activity compared to parental AvPAL*. This is consistent with a recently concluded study on PcPAL, where bulkiness of the residues lining the active site were considered important^15^. F363, K419, and E448 all showed negative fitness for all substituted amino acids sampled and are conserved in PALs and TALs with available structures. I423 showed negative fitness for all mutations, including Thr, the equivalent of which in *Petroselinum crispum* PAL (PcPAL) (I460T) demonstrated a modest 1.15-fold increase in k_cat_^41^. Thus, many of the outcomes of the analysis of DMS data related to active site residues are largely consistent with published data on PALs, supporting the validity of the workflow and calculated fitness scores as proxy for enzyme activity (**Fig. S4**). Further, we identified 4 distinct active site mutations at L108 and M222 that increase AvPAL* activity, only one of which has been previously reported (M222L, by our group)^17^.

### Sequence-function characterization highlights hotspots that enhance activity

To identify “hotspots” that contribute highly to activity across the three passages, we calculated the fitness of each mutation for each passage and then used linear regression to determine the rate of change of fitness over these passages – i.e., their fitness gradient (**Fig. 3a**). We omitted the N-terminal residues since they were previously shown to be dispensable for AvPAL* activity^8^, as well as any mutation with a frequency of zero in any passages or less than 0.625 % in passage #3, to better identify the most significant mutations. From the 225 positively fit mutations found in the passage #3, we identified 12 positions (and 14 mutations) with positive fitness gradients and mapped the fitness onto the structure of AvPAL* (PDB ID: 3CZO) to investigate where the most fit positions are located relative to the active site and one another (**Fig. 3b-e**). Interestingly, we found that the most fit mutations were clustered at either end of the protein chain, with many distal from the active site. We docked phenylalanine (Phe) into the crystal structure of the AvPAL* (2NYN) and calculated the distance of the α-carbon of each residue to the α-carbon of Phe docked in the active site of the same chain (**Fig. 3c**). We saw that 7 of 12 of the fittest mutations were >50 Å from the docked Phe. Noting that AvPAL* is a homotetramer composed of dimers with chains oriented anti-parallel, we found that intramolecular distal residues were proximal to adjacent active sites (**Fig. 3d**). We see that positions 268, 294, 306, 400, and 494 from the B-chain are actually closest to the A-chain substrate Phe, as are the 407, 533 and 534 positions from the C-chain. In fact, previous investigations into the structure of AvPAL* have identified loops in the adjacent chains that play an important role in forming the active site pocket (**Fig. 3c, e**)^8^. Since G218S and A, M222L and N453S had the most abundance in the enriched library, we also investigated if these mutations occur together in any combinations (**Table S2**). We observed that single site mutants were the most abundant in the library whereas triple mutants were completely absent.

**Fig. 3:**
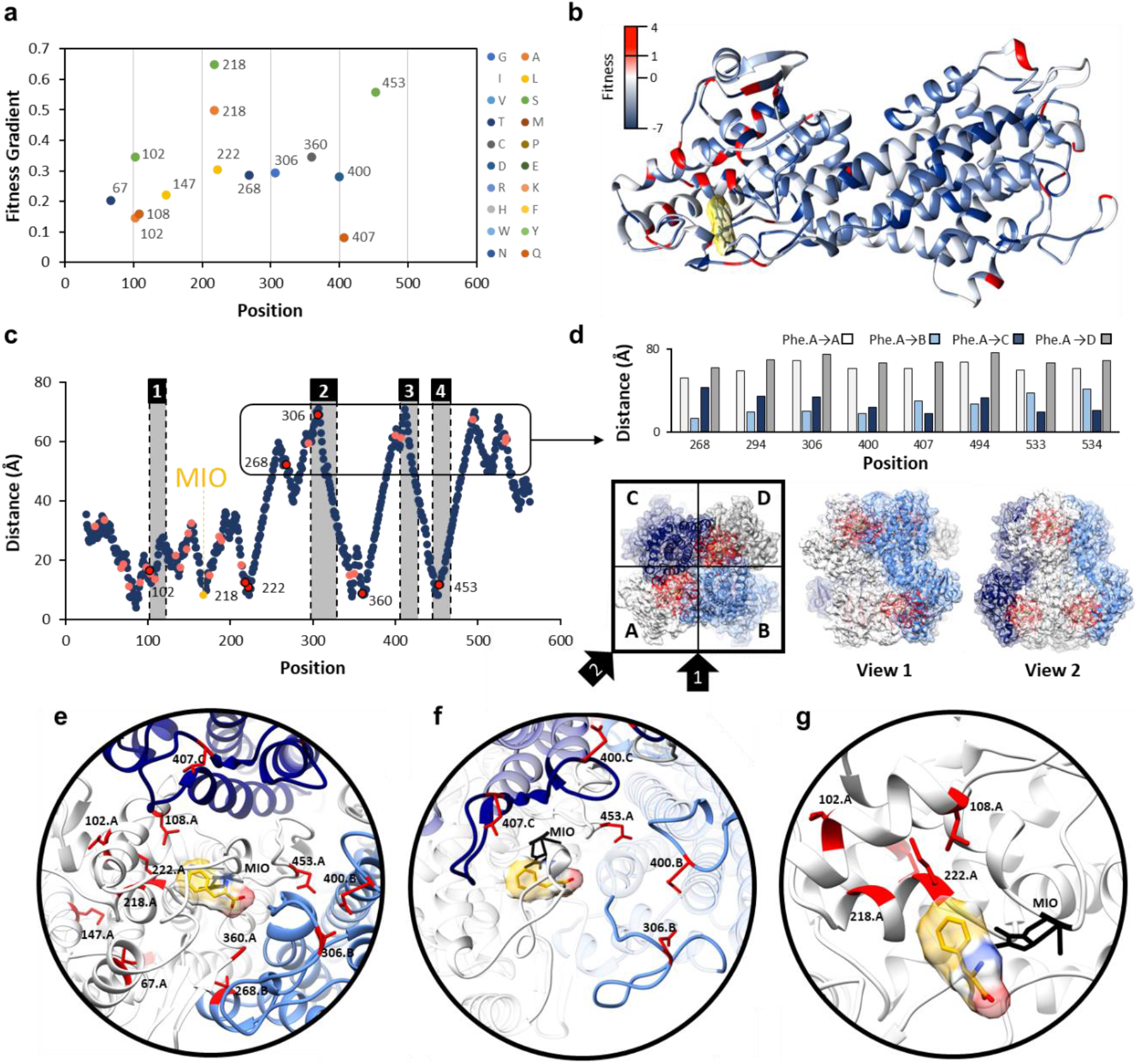
Identification and location of highest fitness positions. **a)** Gradient of fitness is calculated from passages 1-3. Only mutations with a frequency greater than zero in all passages and a passage #3 frequency greater than 0.625 % are shown. A positive gradient indicates increasing fitness across passages. **b)** Fitness values mapped to the structure of a single chain of AvPAL*. **c)** Distance of the α-carbon of docked phenylalanine to the α-carbon of every residue in AvPAL* within the same chain. Residues with a passage #3 fitness < 1 are dark blue, 1−2 are pink, and > 2 are red outlined in black. Active site loops are numbered 1−4 and shaded grey. **d)** High fitness residues that are distal from the active site are proximal to active sites of other subunits when visualized as part of a homotetramer. Chains A (white), B (light blue), C (dark blue), and D (grey) are shown from top and two side views. Residues near the active site are red. **e)** Locations of most fit residues relative to the active site of Chain A. Chains A (white), B (light blue), C (dark blue), and D (grey) are shown. The MIO adduct is black and the phenylalanine in yellow with red and blue oxygen and nitrogen atoms, respectively. Fit residues are colored red with sidechains shown. **f)** Residues 400, 407, 306, and 453 are in the active site loops previously identified as important for active site stability. **g)** Residues 102, 108, 218, and 222, are part of α-helices that form a surface within the active site, near the phenyl ring of the docked phenylalanine.

We next sought to identify mutations at hotpots residues that increase enzyme activity and assess whether global sequence-function characterization can aid in semi-rational design of a highly active AvPAL* variants. The 7 most fit positions selected were 102, 218, 222, 268, 306, 360, and 453 (**Fig. 3e**). We generally classified these residues as comprising 2 regions: and i) residues located within the loops surrounding the active site (268, 306, 360, and 453) (**Fig. 3f**), and ii) residues clustered within a bundle of α-helices that form a surface in the active site (102, 218, 222) (**Fig. 3g**). The 102, 218, 222, 360, and 453 positions most likely act on the intra-chain active site, while the 268 and 306 positions are from an adjacent chain (chain B, relative to the active site of chain A, and vice versa).

To investigate how different mutations interact with each other and shape the fitness, we analyzed the 7 hotspots using single (^7^C_1_ – i.e., 7 choose 1), triple (^7^C_3_ – 7 choose 3), and hepta-site (^7^C_7_ – 7 choose 7) saturation mutagenesis libraries. We constructed the three libraries in a manner allowing us to extensively cover all the combinations practically possible (See material and methods). We then enriched these libraries using our HTS and calculated the fitness of each residue. **Fig. 4a** shows the fitness of single site saturation mutagenesis of all sites when passaged individually (^7^C_1_). For all sites, we found substitutions that were fitter than the wildtype. For instance, at T102, mutation to Ala, Asp, Pro, or His improves fitness. In fact, this position shows the most permissive behavior with seven amino acid substitutions showing positive fitness. The other six positions display more restrictive pattern with only 3−4 amino acids showing positive fitness (**Fig. 4a**). In total we found 28 individual substitutions with higher fitness than the native residue at that position and 24 substitutions with positive fitness. Interestingly, only 2 of these 24 have previously been described to enhance AvPAL* activity (our previous work^17^) – the remaining being new to this work. However, fewer mutations showed positive fitness scores when evaluated in combination using ^7^C_3_ and ^7^C_7_ libraries (**Fig. 4b, c**). This indicates that while many mutations contribute to positive fitness individually, most become negatively fit in combination, likely due to negative epistasis and increased selective pressure due to the presence of more active variants. This agrees with the observation that double mutants from a G218X-M222X library are less fit than single mutants at either of those positions (**Fig. S6**). Indeed, the G218A mutations while highly fit in isolation, is generally unable to contribute significant fitness score when combined with 1 to 6 other mutations (**Fig. 4c, Fig. S6**). We also looked at the naturally occurring diversity at the seven sites investigated in the present study (**Fig. 4d**). For four of the seven positions (218, 222, 306, 360), we find that the natural diversity fully recapitulated the fittest variants found in our study. However, for other sites (102, 268, 453), we were able to find novel mutations that are not predictable through sequence alignment alone. We also noticed that position 306 has very high natural diversity but is restricted to only two mutations with positive fitness (Glu, Gly). Conversely, 453 is naturally restricted to only two amino acids naturally (Asn, Glu) but many alternate mutations increase fitness.

**Fig. 4.**
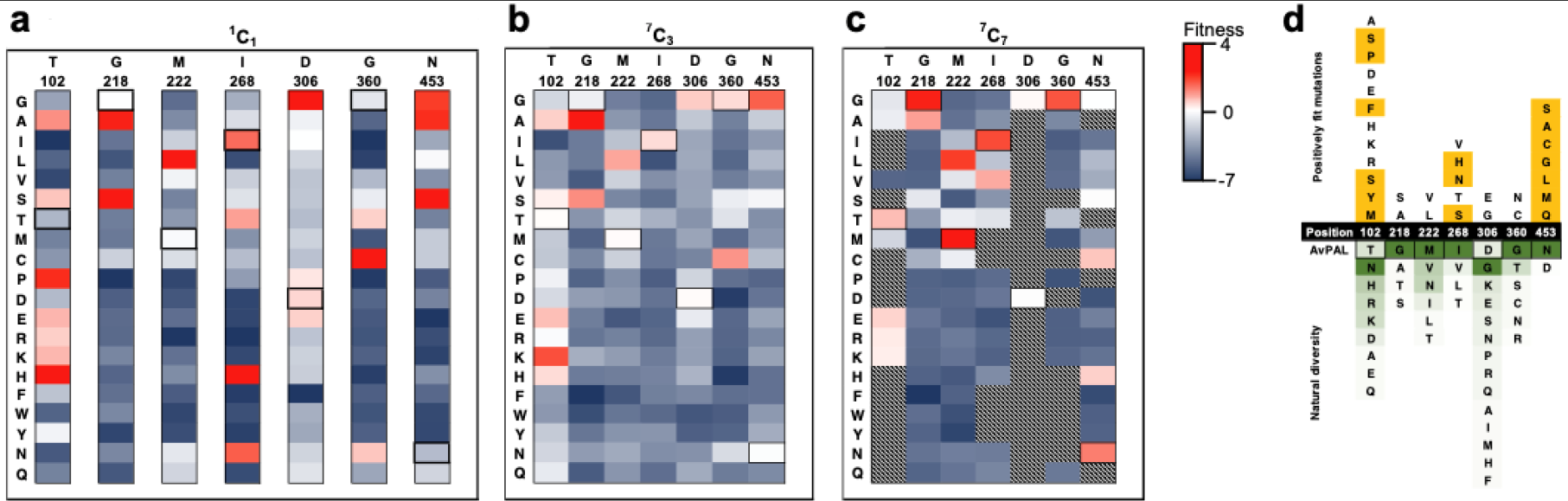
Characterization of site saturation mutant libraries. Fitness heatmaps of **a)** ^7^C_1_, **b)** ^7^C_3_, and **c)** ^7^C_7_ mutant libraries. Relative fitness is shown as two gradients, from highest fitness (red) to zero fitness (white) and zero fitness to lowest fitness (dark blue). Relative fitness for ^1^C_1_ library was calculated at each position individually, and at all positions in combination for ^7^C_3_ and ^7^C_7_. **d)** Sequence comparison to 100 proteins with greatest homology to AvPAL*. Black cells are the 7 hotspot positions, numbered for AvPAL*. Yellow cells indicate positively fit mutations found during our enrichments that were unique relative to the homologous proteins. A green gradient was applied to the natural residues indicating frequency of residue at each position, with dark green as the most frequent and light green as infrequent.

To identify the most active variants from the ^7^C_3_ and ^7^C_7_ libraries, we picked 20 random colonies from the enriched culture and tested for growth and tCA production (**Fig. S7−8**). Upon sequencing variants that showed the highest tCA production, we found mutants with significantly higher activity than parental enzyme (**Table 1**). Among these, T102E-M222L and T102R-M222L-D306G displayed 2.4- and 2.25-fold improvement in the k_cat_, respectively. Interestingly, T102R-M222L-D306G showed some substrate inhibition (**Fig. S9**, apparent at > 10 mM Phe), although, it did display the highest activity at lower substrate concentrations (< 300 μM), which is most relevant for PKU treatment. Other than having isolated the most active AvPAL* variants, these results have two additional and major implications. First, only a small combination of mutations at these sites synergistically and/or additively enhance AvPAL* activity (**Fig 4d**). Second, the propensity of these mutations to act additively and/or synergistically may be explained by the mechanism by which they contribute to PAL activity. To understand the mechanism by which the different mutations may contribute increasing activity, we performed in silico modelling studies.

**Table 1.** Kinetic constants of highest activity AvPAL* variants.

| PAL-variant | Model | $V_{\max}$ | $K_M$ ($\mu\text{M}$) | $K_i$ (mM) | $k_{\text{cat}}$ | $k_{\text{cat}}/K_M$ | Fold increase<br>$k_{\text{cat}}$ |
| --- | --- | --- | --- | --- | --- | --- | --- |
| | | ($\mu\text{mole}\cdot\text{min}^{-1}\cdot\text{mg}^{-1}$) | | | ($\text{s}^{-1}$) | ($\text{s}^{-1}\cdot\mu\text{M}^{-1}$) | |
| AvPAL* | MM | $0.93 \pm 0.01$ | $137 \pm 0$ | - | 0.97 | 0.007 | 1.00 |
| M222L | MM | $1.85 \pm 0.02$ | $145 \pm 7$ | - | 1.93 | 0.013 | 1.99 |
| L4P-G218S | SI | $1.86 \pm 0.02$ | $199 \pm 9$ | $208 \pm 39$ | 1.94 | 0.010 | 2.00 |
| G218S | SI | $1.49 \pm 0.02$ | $180 \pm 9$ | $164 \pm 26$ | 1.55 | 0.009 | 1.60 |
| G218A | SI | $1.01 \pm 0.02$ | $43 \pm 4$ | $96 \pm 18$ | 1.05 | 0.024 | 1.09 |
| T102P | MM | $1.88 \pm 0.02$ | $253 \pm 20$ | - | 1.96 | 0.008 | 2.02 |
| T102E-M222L | MM | $2.33 \pm 0.02$ | $144 \pm 6$ | - | 2.43 | 0.017 | 2.51 |
| T102S-M222L | SI | $1.88 \pm 0.02$ | $111 \pm 6$ | $371 \pm 136$ | 1.96 | 0.018 | 2.02 |
| T102M-M222L-D306G-N453G | SI | $1.15 \pm 0.01$ | $48 \pm 3$ | $147 \pm 24$ | 1.20 | 0.025 | 1.24 |
| T102R-M222L-D306G | SI | $2.16 \pm 0.02$ | $96 \pm 4$ | $169 \pm 21$ | 2.25 | 0.023 | 2.32 |

MM – Michaelis-Menten (Eqn S1), SI – Substrate inhibition, (Eqn S2)

### MD studies reveal mutants show local fluctuations in the active site that impacts near attack conformation

Having identified mutations that enhance AvPAL* activity, we were interested in understanding the mechanism contributing to increased activity. We conducted extensive all-atom atomistic MD studies for different mutants of AvPAL* to ascertain the stability of attack conformation. The starting enzyme substrate (E-S) complex was derived using docking studies; each MD simulation was 500 ns × 2 long; and all post-simulation studies were conducted from 100^th^ ns onwards. AvPAL* is a homotetramer, of closely interlocking monomers^6,8,36^. Each tetramer contains four catalytic sites, and each active site is comprised of residues from three different monomers and one MIO group. Each active site is capped by two flexible loops; an inner loop which is packed tightly in the active site and forms interactions with the substrate and an outer loop which serves as an external cap. The outer loop is attributed to forming a barrier to bulk solvent preventing access to the active site^8^. Reaction mechanism of PALs have been extensively studied and involves the formation of N-MIO intermediate (**Fig. S10**)^25,30^. Here, we focused on the formation of the first intermediate state of the substrate in the active site (before covalent binding with MIO, **Fig. S10a–b**). During this step, Y314 functions as the catalytic base.

As the binding conformation of the substrate is not yet identified in any of the PAL crystal structure. We performed the docking of Phe in AvPAL* structure using Autodock4 and then chose energetically and structurally feasible conformation for further studies. We observed the binding energy to be −3.03 kcal mol^−1^ for the E-S complex. Maintenance of close proximity between substrate and catalytic residues over the period of the MD simulation is indicative of a stable E-S complex and formation of near-attack conformation. We measured the distances between substrate amino nitrogen (Phe(N)) and the MIO methylidene carbon (MIO(Cβ2)) and adjacent chain tyrosine 314 hydroxyl oxygen (Y314(O)) for the variants (**Fig. S11**).

Hyperactive variants, M222L and G218S, but not N453S, show better near-attack conformation as indicated by closer proximity to MIO when compared to controls, parental AvPAL* and low activity variant L108G (**Fig. 5a−d**). We use L108G as a control for our modeling studies as it has previously been validated as a deleterious mutation^8^. The double and triple mutants, T012E-M222L and T102R-M222L-D306G, also interact closely with the reactive MIO(Cβ2) compared to controls. Next, we calculated root mean square fluctuation (RMSF) for mutant backbone atoms and normalized them over the parental AvPAL* to identify the flexible regions. Higher values are characteristic of flexible regions that readily displace from their average position in the parental enzyme during simulation and vice-versa for negative values. Four flexible regions in the high-activity single mutants that show altered dynamics compared to parental are residues 60−130, 200−230, 280−325, and 440−460 (**Fig. 5g, S11**). These regions are located around the active-site and the interface of the dimeric subunits. The loop residues 285−325 constitutes an access channel of the enzyme and residues 310−318 forms the second shell of the active site. Regions 60-130 and 440−460 interact with phenyl and carboxylate group of the substrate, respectively. From **Fig. 5g**, we concluded that the dynamics of AvPAL* changed with every mutation and the negative control (L108G) showed very high fluctuation in the access channel and second shell (285−325) that likely disrupt profitable substrate interactions. In addition, we also evaluated the Free Energy Surface (FES) obtained from different sets of metadynamics experiment which provide us an understanding of the energy bins associated within the active site and the regions around it, which further supports the assertion that beneficial mutants stabilize interactions between the substrate and active site (**Fig. S12**). Overall, our MD simulations suggest that for three of the four hyperactive variants (M222L, G218S, T012E-M222L, and T102R-M222L-D306G) the substrate more readily approaches the catalytic site forming a stable near-attack conformation. However, for N453S, our analysis revealed behavior largely unchanged from parental, suggesting that its mode of action may not involve direct modulation of interactions between substrate and catalytic residues.

**Figure 5.**
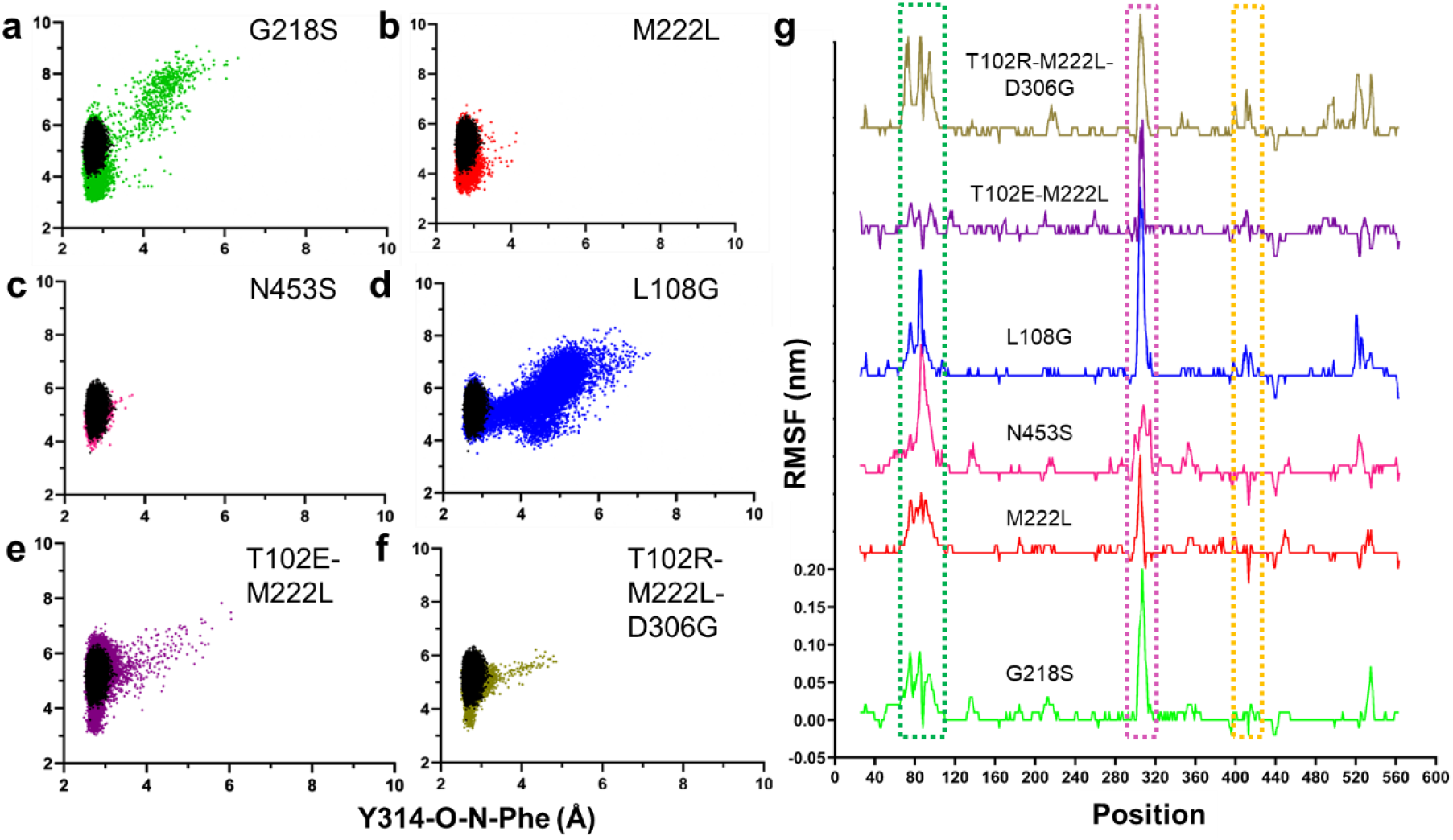
Results from MD. Scatter plots of distance calculated between Y314(O)−Phe(N) and MIO(Cβ2)−Phe(N) atoms in parental AvPAL* (black), **a)** G218S (green), **b)** M222L (red), **c)** N453S (pink), **d)** L108G (blue), **e)** T102E-M222L (purple), and **f)** T102R-M222L-D306G (khaki). **g)** RMSF (root mean square fluctuation) plot of protein backbone atoms (carboxylate, Cα, amine). The RMSF values of mutants are normalized to that of the parental enzyme so that only major movements are amplified. All variants are plotted on the same scale.

### Steered molecular dynamics (SMD) studies show steady and seamless diffusion of Phe in mutant N453S

MD studies with N453S reflected behavior similar to that of parental AvPAL*. As N453S is in the periphery (~12 Å from the active site), we suspected it impacted Phe diffusion. We therefore performed SMD of AvPAL* and the mutant for comparison studies. One of the near attack conformations was chosen to simulate the egress path taken by the Phe from the active site. Subsequently, we used the same path was as a shadow to simulate the Phe reassociation studies. With the primary force constant, both egress and the reassociation of the Phe favored substrate diffusion in N453S compared to AvPAL* (**Fig. 6, Movie S1−S2**), details are explained in the supplementary section (**Fig. S13**). We conducted umbrella sampling as an extension of SMD studies to estimate the energetics during the translocation of the Phe in the path. A series of configuration or reaction coordinates across the path were chosen from the SMD studies and constructed based on the distance between the COM of MIO and that of the Phe. The path was discretized into multiple windows that were chosen for every 0.5 Å of the Phe movement from the active site till it reached the periphery of the protein. The umbrella sampling studies on Phe translocation sheds light on mutation N453S and the residues along the path that are responsible for the substrate stabilization and anchoring as it enters the active site. The potential of mean force (PMF) graph shows that N453S has two minima, which were not observed in the AvPAL*, at a distance of ~5−6 Å and ~10−11 Å between the COM of MIO and that of Phe (**Fig. 6a**). The conformational changes of Phe were extracted from umbrella sampling and mapped on the protein for AvPAL* and N453S (**Fig. 6b-c**). For AvPAL*, the path is narrow towards the active site leading to slightly constrained and energetically less favorable entry (**Fig. 6b**). The path for N453S is wider and energetically favorable, especially in a region close to the active site where the substate shows well organized conformations projecting the amino group towards the positively charged residues (**Fig. 6f**). Due to this, the phenyl group of Phe likely enters the active site and forms a precise Michaelis complex (**Fig. S14**).

**Figure 6.**
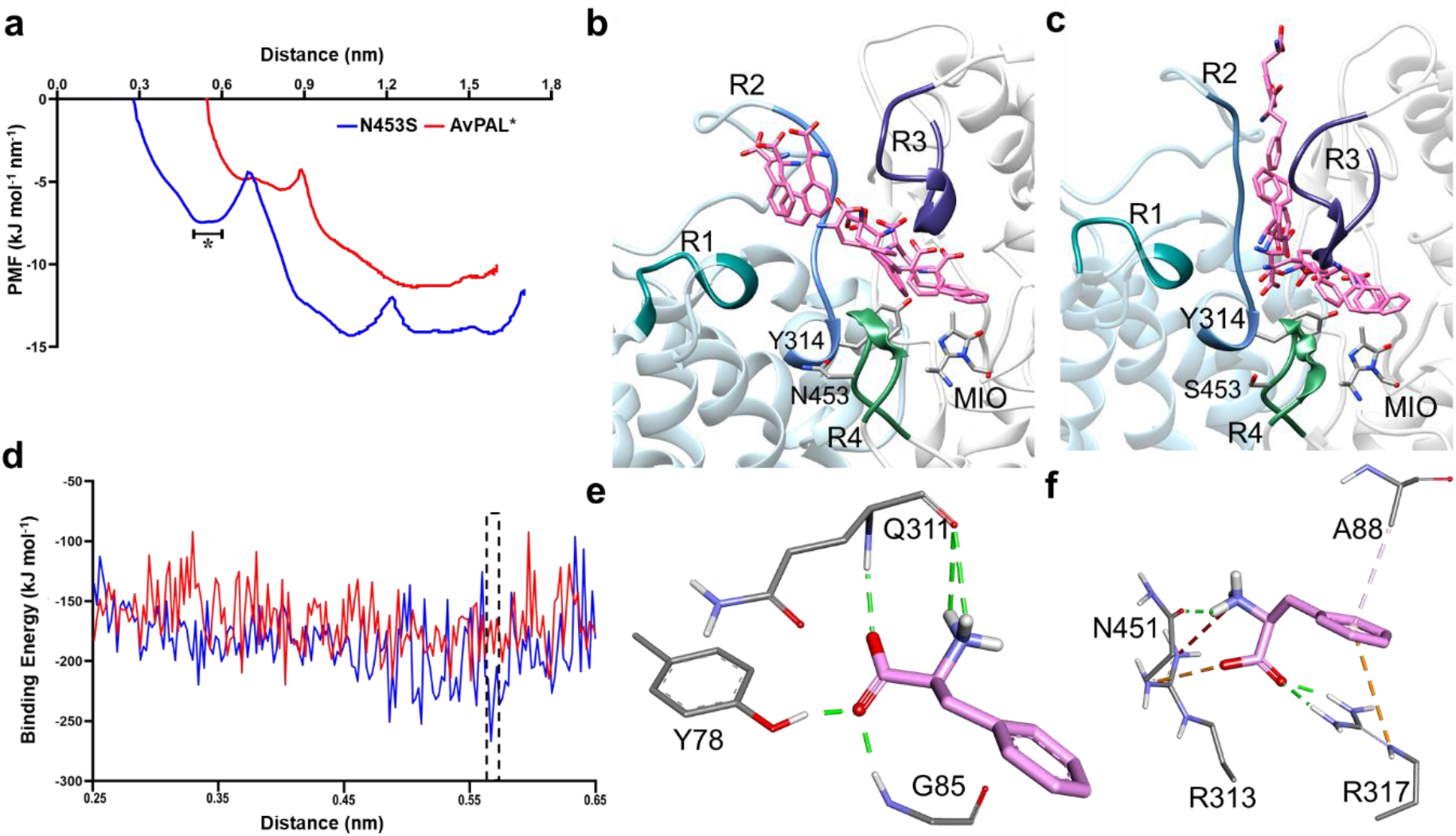
Results from SMD and umbrella sampling for parental AvPAL* and N453S. **a)** The conformational transition of the substrate along the PMF profile, **b)** extracted from the parental and **c)** mutant N453S. The peripheral regions of the substrate entry path are highlighted as R1, R2, R3, and R4. They composed of residues 397−403, 308−315 of chain C and 83−94, 446−455 of chain A, respectively. The region marked in as (*) in (a) is the free energy dip that facilities the substrate entry in N453S. **d)** Binding free energy calculations showed higher affinity of Phe for mutant N453S. **e, f)** E-S complex extracted from free energy calculations with least binding energies for parental and mutant (dotted box in (d)) reveal that the substrate is stabilized by salt bridges and hydrogen bonds in N453S and only hydrogen bonds in the parent. Green lines show hydrogen bond interactions, orange lines are salt bridge interactions, and light pink lines are hydrophobic interactions.

In addition, we calculated the binding energy for E-S complex from region that showed differences in AvPAL* and N453S denoted as asterisk (*) in the PMF profile (**Fig. 6a**). The binding energy for N453S improved by ~35 kcal mol^−1^ when compared to AvPAL* (**Fig. 6d**). The conformation of the substrate and its interacting residues were extracted from the region denoted by asterisk in **Fig. 6b−c** that represents low energy region for both AvPAL* and N453S. In AvPAL*, Phe was found to interact with Y78, the backbone of Q311 and G85, whereas in case of N453S, Phe showed interactions with A88, R313, and R317 (**Fig. 6e−f**). In N453S, A88 was observed to have π-alkyl interaction and R317 shows π-cationic interaction with the phenyl ring of Phe. Among electrostatic interactions, the salt bridge plays a major role in stabilizing and anchoring Phe. R313 and R317 shows salt bridge interaction with the carboxylic group and N451 interacts with the amino group of the Phe. In AvPAL*, Phe amine interacts with the backbone of Q311, the carboxylic group interacts with the backbone nitrogen of G85 and hydroxyl group of Y78 with hydrogen bond interactions (**Fig. 6e**). We did not observe any interactions with phenyl ring that could slow movement of the substrate in either case.

Since N453S showed improved access of the substrate Phe to the active site, we hypothesized that combining it with new variants might further improve the activity. Based on this we generated new combined variants with M222L, L4P-G218S, T102E-M222L and T102R-M222L-D306G. We purified these variants and performed the kinetic characterization (**Fig. 7a−e, Table S3**), determined the whole cell conversion of Phe to tCA (**Fig. 7f**) and growth rate (**Table S4**). We found that all the active N453S added variants displayed similar kinetic parameters as parental AvPAL* (**Fig 7a−e**, **Table S3**), despite their parental counterparts having >2-fold higher activity. Further, presence of N543S in G218S and T102R-M222L-D306G backgrounds completely abolished activity (**Fig. 7c, e**). The reduced vmax for all new variants was surprising because all the active N453S combinations displayed improved whole cell tCA conversion when compared to their parental counterparts (**Fig. 7f**). In fact, T102E-M222L-N453S gave >6-fold higher conversion of Phe to tCA when compared to AvPAL* (**Fig. 7f**). Since the SMD studies suggested more favorable substrate ingress in N453S, we hypothesized that Phe may more readily displace tCA from the active site, reducing product inhibition, a known issue with PALs^42–44^ and may explain higher fitness in cellular context. To test this, we determined the activity of AvPAL*, N453S, T102E-M222L and T102E-M222L-N453S in the presence and absence of 150 μM tCA (**Fig. S15**). We found that in both cases, the N453S variants were less inhibited by tCA. Since T102E-M222L-N453S displayed 6-fold better whole cell conversion of Phe to tCA, it might be a good potential candidate for bacterial- or cell-encapsulated enzyme-replacement therapy for PKU.

**Figure 7.**
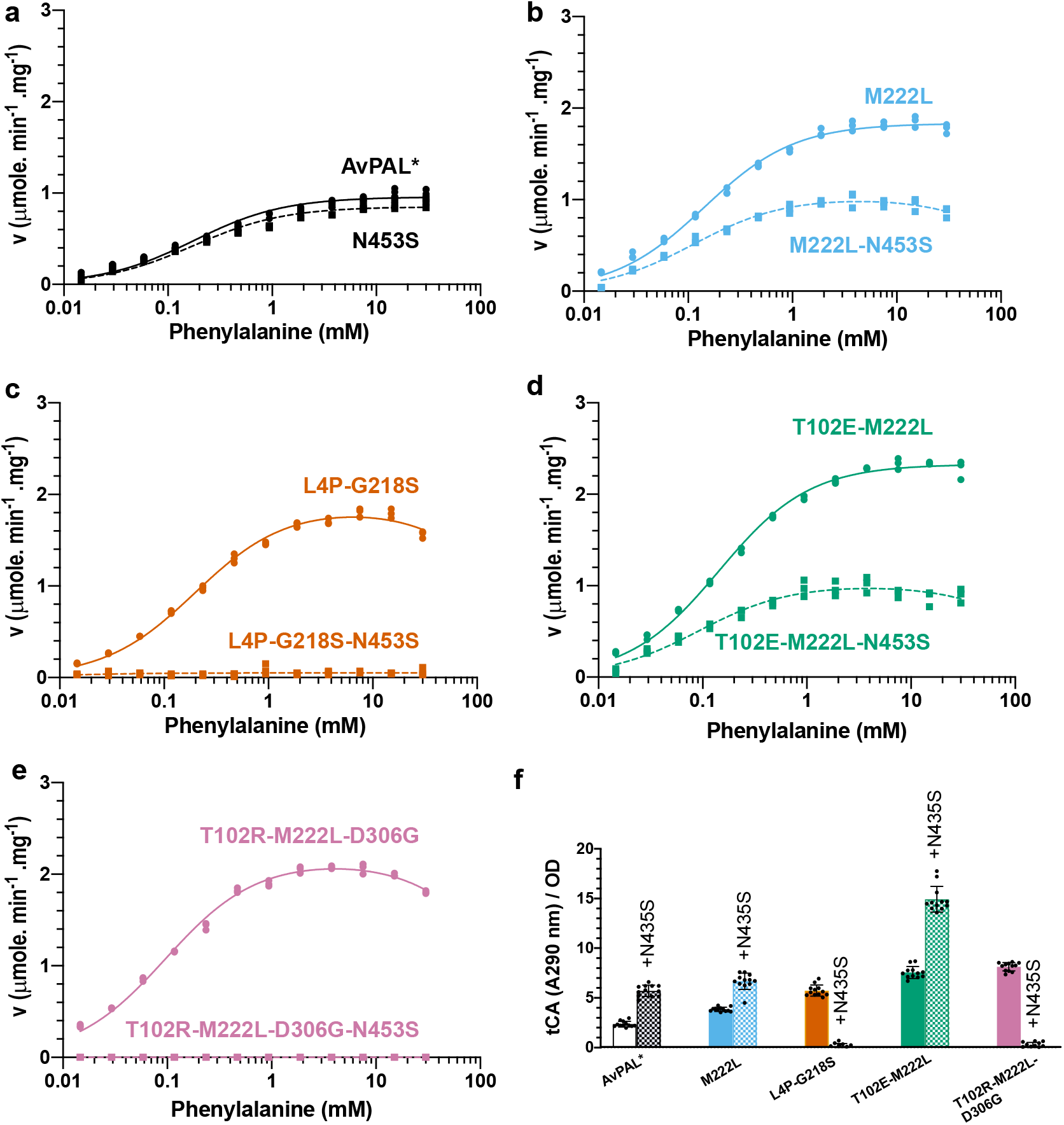
Kinetic characterization and whole cell activity of N453S variants. **a−e)** Michaelis-Menten plots of AvPAL* and high active variants combined with N453S. L4P-G218S-N453S and T102R-M222L-D306G-N453S did not exhibit any activity at any of the Phe concentrations tested. Kinetic parameters are listed in **Table S3**. **f)** Whole cell conversion assay indicates that active enzymes are more active when encapsulated in *E. coli* cells even though there do not display superior kinetic parameters.

### QM/MM reveals stabilization of the transition state in the hyperactive active mutants

Although two reaction mechanisms have been proposed for PALs that proceed either through a Friedel-Crafts (FC) like intermediate^45^ or an N-MIO adduct^30^, there is increasing support for the latter^25^. So, we simulated the N-MIO adduct reaction mechanism that involves formation of near attack conformation where Phe is oriented suitably for proton abstraction by the hydroxyl group of Y314 (**Fig S10a−b**). This deprotonation results in formation of the nucleophilic amino moiety activating Phe for interaction with electrophilic MIO^30^. These rearrangements are referred to as the first step. The starting point of the E-S complex derived from well equilibrated MD simulations (1 μs simulations), shows the least distance between reactive groups, i.e., Y314(O)−Phe(N) and MIO(Cβ2)−Phe(N). To get the least distance between the reactive groups for the QM/MM simulation, we scanned across the MD simulation and this E-S coordinate (MD-ES, **Table S5**) for the distance between the combined center of mass (COM) of MIO and Y314 and that of the amino group of the substrate.

All forms of quantum chemical calculations performed in this study were to delineate the reaction pathway and free-energy barrier for first step of proton abstraction and the intermediate state 1 (IS1) formation where the substrate amine forms an attack conformation with MIO(Cβ2) (**Fig. 8**). First step of the investigation based on the generally proposed mechanism as shown in **Fig. S16**. We obtained the detailed reaction mechanism, the optimized structures of transition states and intermediates which are shown in **Fig. 8a-c** and the calculated overall relative free energy graph is given in **Fig. 8d**. To understand the mechanism, we implemented the new functionality by NAMD that can execute multiple QM regions in parallel^46^. All four active sites of PAL were treated under QM code simultaneously while the rest of the protein was treated under MM force field (**Fig. S17**). The E-S complex is defined as the zero-point Ground state (GS, 0 kcal mol^−1^) to which all other energies are compared for each mutation. To define the attack conformation from Michaelis complex, we conducted QM/MM using PM7 function by including active site residues Y314, Q452, R317, MIO, and Phe until the distance between the Y314(O) and substrate amine approached ~1.5 Å (**Table S6**). The distance we observed is similar to that derived by QM/MM on TAL from *Rhodobacter capsulatus*^47^ and crystal structure from PcPAL^25^. This was used as starting point reaction coordinates for transition state (TS) optimization using higher level QM/MM simulations based on B3LYP def2-SVP D4 TS. TS and IS were studied till the substrate H^+^ were abstracted by Y314, and then until the Phe moved to an optimum attack conformation with MIO. We observed the following sequence of events in the first step of the reaction in parental AvPAL*. After the near-attack conformation was formed, the bond between the Y314(O) and abstracted H^+^ was stretched from 1.04 Å to 1.21 Å and at the same time, the carboxyl oxygen of the Phe abstracts the H^+^ from Y314. First transition state (TS1) is formed when the H^+^ from the amino nitrogen is stretched to 1.28 Å, before being completely extracted by Y314, followed by a small conformational change in the Phe that brings it closer to MIO to form IS1. For all the tested mutations same events were observed for the first step but the mutations showed different energies for the TS1 and IS1. The energies for both the TS1 and IS1 correlated with the experimental data and the energies can be graded as L108G < AvPAL* < M222L < G218S< T102E-M222L < T102R-M222L-D306G and suggest that formation of IS1 through TS1 may be rate-limiting in AvPAL* but shifts to a downstream step in all mutants other than N453S (**Fig. S18**). The conformational changes observed in during the formation of the IS1 is described in supplementary information and **Fig. S19**.

**Figure 8.**
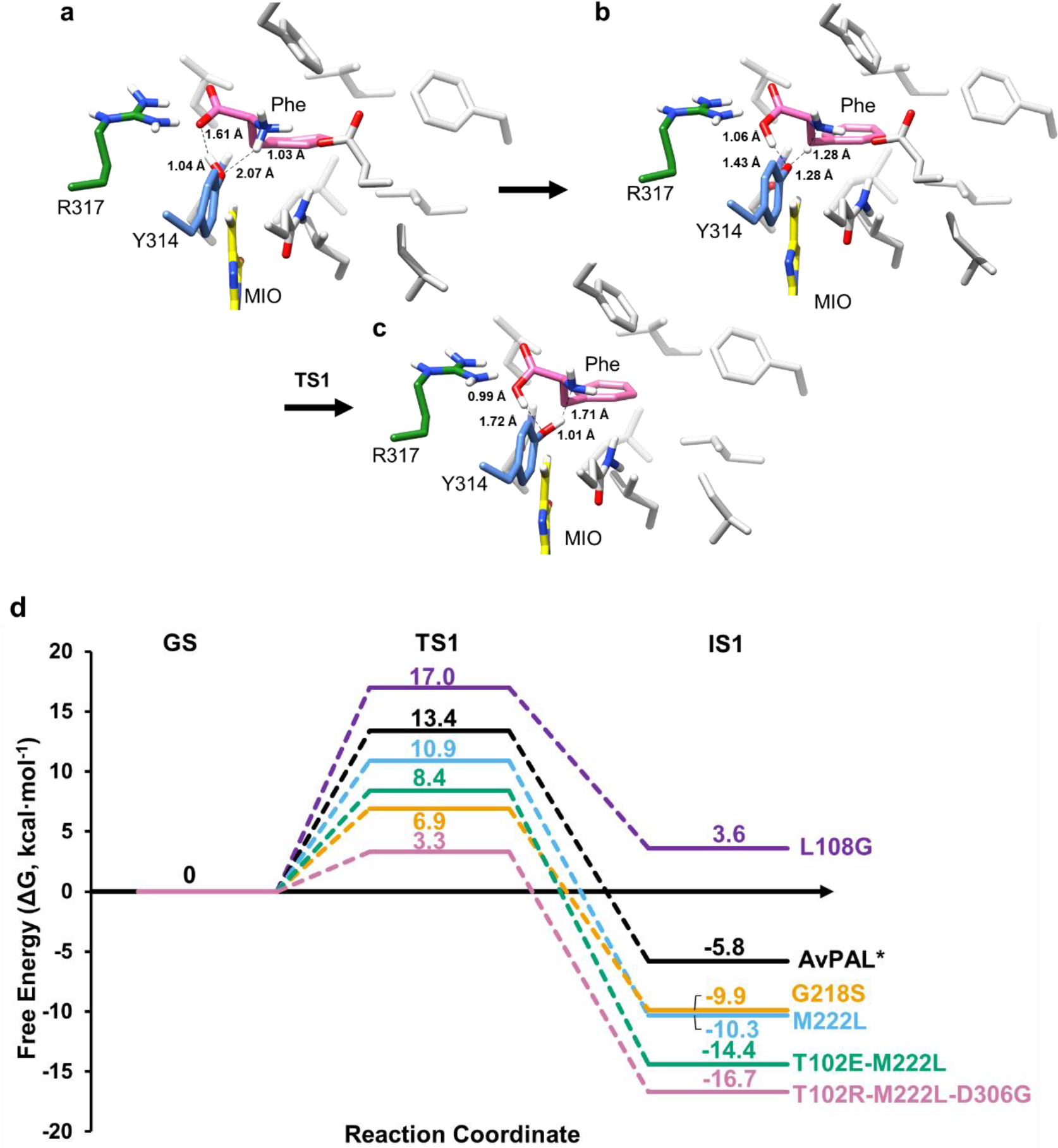
QM/MM studies on AvPAL* variants. Proposed reaction mechanism. **a-c)** Transition states for the first step of the reaction, where proton transfer takes place, with Phe (pink), MIO (yellow), catalytic Y314 (blue), and R317 (green), and other protein residues (light grey) as stick models. Only polar hydrogens are shown for clarity. All distances are in Å. **d)** Relative energy landscape for AvPAL* (black), M222L (red), G218S (blue), L108G (green), T102E-M222L (pink), and T102R-M222L-D306G (cyan) for all the steps from ground state (GS) to IS1. Energies along paths are not to scale.

Next, to understand the barrier-crossing events shown in the QM/MM simulations in depth, we conducted hybrid QM/MM techniques combined with metadynamics, which enhances the sampling of coordinates relevant to the reaction. This way we can observe how the system accelerates across the reaction barriers by itself and escapes from local minima (**Fig. S20**). We further characterized the TS1 and IS1 structures by transition path sampling (TPS) simulations, and this was plotted over the free energy surface (FES). FES and TPS derived from QM/MM metadynamics could clearly differentiate the mutants and AvPAL* and showed thermodynamically favorable energy paths for T102E-M222L and T102R-M222L-D306G (details in supplemental information, **Fig. S20**).

## CONCLUSIONS

In summary, we report the most extensive sequence-function analysis of an MIO-containing enzyme, AvPAL*, that leveraged a growth-coupled HTS. The outcomes of DMS guided identification of mutants that enhance native PAL activity. Further, by performing computational studies (QM/MM, MD, steered MD + QMM/MM), we identified the mechanisms through which the mutations enhanced enzyme activity, which in turn, allowed us to identify variants that have promising applications to cell-free biocatalysts (T102E-M222L, T102R-M222L-D306G) and cell-based systems (T102R-M222L-N453S). Not only does this significantly advance enzymology and engineering of PALs, but also demonstrates the power of using DMS to guide basic and applied enzymology.

## METHODS

### Strains and general techniques for DNA manipulation

AvPAL* error prone library enriched previously was used in the current study^17^. PCR was performed using Phusion DNA polymerase or Platinum™ SuperFi II Green PCR Master Mix (ThermoFisher Scientific). *E. coli* NEB5α (New England Biolabs) was used for plasmid propagation and *E. coli* MG1655 *rph+* was used for screening of libraries and purification of recombinant AvPAL* and its mutants. Sequences of constructed plasmids were confirmed through DNA sequencing (Genewiz). AvPAL* was expressed under constitutive T5 promoter from plasmid pBAV1k carrying chloramphenicol resistance.

### Next generation sequencing (NGS) of library and data processing

Plasmid libraries and PCR products were outsourced to Genewiz (New Jersey, USA) for sequencing on Illumina MiSeq Nextra paired end sequencing platform (2 × 250 bp). For all the samples sequenced we received 1−5 million reads with average length of 160 bp after trimming. The bioinformatic workflow is depicted in **Fig. S21**. Briefly, the raw fastQ files were evaluated for quality score, read length, adaptor and duplicate read content using FastQC package. Subsequent analysis was performed using Geneious Prime® 2020.2.4. The reads were paired and merged using BBMerge package^48^, filtered for adaptor sequences, short and poor-quality reads using BBDuk package. The reads were then mapped onto reference gene (AvPAL*) using BowTie2 package^49^. The mapped reads were then analyzed for single nucleotide variants to detect mutations. This variant call file was used to calculate the fitness score using **Eqn 1**.

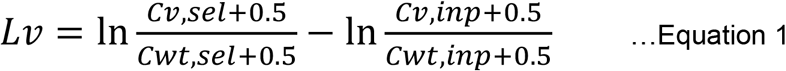

The fitness values thus determined were represented using heatmap to show the residues with positive and negative fitness.

### Physical linking of distal mutations for amplicon sequencing

Workflow followed for physical linkage of G218, M222 and N453 is indicated in **Fig. S22**. Briefly, region immediate downstream to M222 and immediate upstream to N453 was amplified with primers having homologous region. The amplicon flanked by homologous region was sealed using NEB-HiFi assembler. The circular plasmid was then used as a template to amplify ~300 bp region spanning G218-N453. The segment was amplified using primers with illumine sequencing overhangs. The amplicon was sequenced using AmpliconEZ Seq Illumina platform at Genewiz.

### Construction of site saturation mutagenesis (SSM) libraries

pBAV1k plasmid containing AvPAL* was used as template for constructing the site saturation libraries. The SSM libraries for seven sites of interest were constructed in three ways; i) individual sites using NNS codon at the target location was constructed following QuickChange-like method. Briefly, partially overlapping primers were used to perform inverse PCR, the amplicon was subjected to DpnI (NEB) digestion to remove the parental plasmid followed by NEBHiFi assembly (NEB) to assemble and seal the overlapping ends for improved transformation efficiency. The assembled product was purified using PCR clean-up kit and electroporated into MG1655 *rph+.* ii) A new approach of scaling by mutation was developed to mutate three sites in varying combination. In this approach, the clean-up product from approach i) was pooled in equimolar amounts and used as template for second round of inverse PCR using seven primer pairs individually. This process was repeated total of three times to generate the ^7^C_3_ SSM library which was transformed into *E. cloni* DH10B (Lucigen) for achieving large library size. iii) The third library was constructed by using restricted codon at the seven sites of interest – ^7^C_7_. The restricted codon was chosen based on the DMS data from error prone PCR library screen (**Table S7**). The fragments were assembled using NEB HiFi assembler and electroporated into *E. cloni* DH10B after PCR clean-up. The plasmid library from approach ii) and iii) were isolated from *E. cloni* and transformed into *E. coli* MG1655 *rph+* enrichment on minimal media containing 30 mM Phe. Fitness data for these libraries were obtained by sequencing the seven sites of interest using AmpliconEZ seq (Genewiz). The data was processed as described in a manner described above.

### Enzyme assay, purification, and kinetic characterization

PAL activity was monitored by measuring the production of tCA at 290 nm over time. Briefly, 200 μL reaction as performed by 1 μg of purified enzyme to pre-warmed PBS containing 30 mM Phe. The assay was performed in 96-well F-bottom UVStar (Greiner Bio-One, Kremsmünster, Austria) microtiter plate and absorbance at 290 nm was measured every 15 s at 37 °C using a SpectraMax M3 (Molecular Devices) plate reader.

For purification, the enzyme was isolated from 25 mL culture. The pellet was washed once with PBS and resuspended in 500 μL PBS. This cell suspension was sonicated on ice using a Sonifier SFX 150 (Branson Ultrasonics, Danbury, CT) (10 s ON; 1 min OFF; 2 min; 40 %), and cell debris was separated from the lysate by centrifuging at 20,000 × g for 10 min at 4°C. As each construct included a N-term His-tag, the enzyme was purified via immobilized metal affinity chromatography (IMAC) purification. Briefly, the lysate was loaded onto HisPur™ Ni-NTA Spin Plates (ThermoFisher Scientific) and incubated for 2 min. After being washed four times equilibration buffer, pure protein was then eluted using 200 μL of Elution buffer (300 mM NaCl, 50 mM NaH2PO4, 500 mM imidazole, pH 8.0). Elution fractions were then dialyzed using Tube-O-Dialyser tubes (1 kDa MWCO, Geno-Tech). Protein concentration was estimated by Bradford reagent (VWR) using bovine serum albumin (BSA) as the standard. For kinetic analysis, AvPAL* and selected mutants were purified and assayed as described above. The activity was measured at twelve concentrations of Phe ranging from 15 μM to 30 mM in PBS, pH 7.4 (PBS) at 37 °C. A Michaelis-Menten curve was fit in GraphPad Prism software using the initial rate at each Phe concentration.

### Modelling and induced-fit conformation sampling/enzyme-substrate interaction studies

3D structure of the PAL enzyme from *Anabaena variabilis* chosen (PDB ID: 2NYN). The structure had 2 missing regions (Residues 74-92, 302-309) which were modelled using MODELLER^50^. The PAL structure with the least DOPE Score was selected and chosen for further studies. The binding conformation of phenylalanine is not identified in any AvPAL* crystal structures. To compensate, Phe was docked in the active site of the modelled AvPAL* structure using Autodock4 tool^51^. An energetically and structurally feasible conformation was chosen for the interaction studies and structural analysis in Chimera. The binding energy was found to be −3.03 kcal mol^−1^ for the E-S complex.

### Molecular dynamics (MD) simulations

MD simulation was conducted for AvPAL* and mutant complexes from the interaction studies. The complexes were taken into a system using the AMBER99SB^52^ force field as implemented in GROMACS^53–57^ tools. The complex was placed in a box of volume 1000 nm^3^ and then solvated with ~26,230 water molecules. To emulate conditions similar to in vitro experiments, a salt concentration of 0.15 M NaCl was incorporated into the solvated system. This has the added benefit of neutralizing the charge of the system. The LINCS was employed to constrain bond length and fix all bonds containing hydrogen atoms. Berendsen thermostat^58^ was chosen to control the temperature at 310 K. The Particle-mesh Ewald algorithm (PME)^59^ was used to calculate electrostatic interactions with a 10 Å cut-off. The V-rescale and the Parrinello– Rahman algorithms was applied to couple the temperature and pressure. Energy minimization of the system was obtained using the steepest descent algorithm with a tolerance value of 1000 kJ mol^−1^ nm^−1^ in 1000 steps. The minimized system was equilibrated for 1 ns each of constant volume and constant pressure ensemble. The system was then subject to a production run of 500 ns at 1 atm pressure and 310 K, twice for statistical significance. The coordinates obtained from the production run were used for post-simulation analysis to observe the effect of the mutations on the dynamics of the protein. The distances between MIO methylidene atom and substrate amino nitrogen (MIO(Cβ2)−Phe(N)) and Y314 hydroxyl oxygen and substrate amino nitrogen (Y314(O)−Phe(N)) were considered and plot against each other as a scatter plot. Backbone atoms of domains around the active site were considered to calculate Root mean square fluctuation (RMSF).

### Metadynamics-based MD simulations

Metadynamics simulations were performed to understand the free energy landscape of the active site. On completion of simulations, the substrate was expected to find different potential minima to attain near attack conformation in the active site. Comparative studies were conducted for AvPAL* and mutations. In metadynamics based approaches, the choice of a collective variable (CV) in the design of the experiment is crucial. We chose two CVs i.e., distance between COM (center of mass) of substrate atoms and COM of Y314 atoms (CV1), and distance between COM of substrate atoms and COM of heavy atoms in the backbone of residues in conserved secondary structures that were present within 5 Å of G218, M222 and L108 residues (CV2) (**Fig. S12a**).

### Steered molecular dynamics (SMD) studies

SMD simulations were conducted to identify conformational changes and associated path samplings when the substrate is exposed to mechanical strain or rupture force, which cannot be achieved through standard MD simulations. Well equilibrated systems were chosen to be starting points for the SMD studies. The pulling simulations were implemented using GROMACS tools. Substrate was pulled by its COM away from the active site and pulled towards COM of MIO group in unbinding and reassociation process, respectively. The pull velocity of 0.0005 nm^−1^ ps^−1^ with the bias force constant of 310 kJ mol^−1^nm^−2^ and −30 kJ mol^−1^ nm^−2^ were used in unbinding and entry process, respectively. Umbrella sampling were conducted as an extension of SMD studies to estimate the energetics during the translocation of the substrate in the path. A series of configuration or reaction coordinates across the path were chosen from the SMD studies and constructed based on the distance between the COM of MIO and that of the substrate. The path was discretized into multiple windows which were chosen for every 0.5 Å of the substrate movement from the active site till it reaches the periphery of the protein.

### QM/MM simulation

E-S complexes were placed in a cubic box with a solute-solvent separation margin of 12 Å in each dimension, by means of QwikMD^60^ program implemented in VMD. The electroneutrality of the system was maintained by the adding NaCl to maintain a salt concentration of 0.15 M. CHARMM36 forcefield was used for the protein topology (generated using psfgen and autopsf programs) and TIP3P water models were used in the system. During the simulations, a 12.0 Å cut-off was applied to short-range, non-bonded interactions, whereas long-range electrostatic interactions are treated using the particle-mesh Ewald (PME) method. The equations of motions were integrated using the r-RESPA multiple time step scheme to update the short-range interactions every step and long-range interactions every two steps. The time step of integration was set to be 2 fs for all simulations performed. Thermal equilibrations were conducted by first subjecting the system to energy minimization using the conjugated gradients method for 1000 steps (2 ps) and then coupled with a heat bath kept constant at 300 K by the Langevin thermostat with a collision coefficient of 1 ps^−1^ and a barostat maintained at 1 atm.

The last step of classical equilibrium was taken to QM/MM interface to select QM region and initiate QM/MM simulation using QwikMD interface provided in VMD. Four regions in different chains constituting MIO adduct, Q452, Y314, substrate and water molecules within 3.5 Å of the MIO were selected as QM regions, the total charge for each QM region was maintained between +1 and −1 for effective Semi-Empirical QM Calculations.

The system was optimized by a 1,000 steps minimization, followed by 10,000 steps of simulated annealing calculation, equilibration and subjected to an average of 5000 ps QM/MM hybrid production run using PM7^61^ together with the CHARMM36 force field. The least distance between the reactive groups were considered as the good guess of the transition state geometry. This was followed by an average of 1000 ps QM/MM hybrid production run using PM7-TS. Simultaneously, an average of 1000 ps density function theory (DFT) based QM/MM hybrid production at B3LYP/ 6–31G(d) level def2-SVP level implemented in ORCA^62^ was carried out and NEB-TS was used to find and optimize TS. Three input files were provided for the purposes of the QM/MM hybrid production run i.e., i) initial conformation from well equilibrated MD, ii) the transition state geometry derived form QMMM-PM7 and the final product (refer IS1 in **Fig. 8**) that was manually modelled using the transition state geometry, equilibrated using QMMM-PM7 simulation.

The mutants and the AvPAL* were subjected to QM/MM simulation protocol. In this report only the first step of the reaction and their corresponding energy values of PM7-TS are reported. DFT calculations were used only for validating the reaction coordinates of the transition states and the intermediate state.

### QM/MM Metadynamics

5000 ps equilibrated QM/MM reaction coordinate of the E-S was used as initial input structure for the metadynamics simulations. QM/MM-metadynamics simulations was carried out at 300 K, 1 bar, 0.5 fs time step and periodic boundary conditions for 1000 ps using NAMD 2.13^46^ and colvars module^63^. The distance between amino-nitrogen of the substrate & hydroxyl-oxygen of Y314 and distance between amino-nitrogen of the substrate and methylidene carbon of MIO adduct were used as two collective variables (CVs). Gaussians of height 0.2 kcal/mol were added onto the CV coordinate at every step to construct metadynamics bias potential with width of 1°. QM/MM hybrid production run using PM7-TS integrator.

## DATA AVAILABILITY

Deep sequencing data has been submitted to NCBI SRA and is available under accession # PRJNA730338.

## CODE AVAILABILITY

All open-source and commercial software used are described in the methods section.

## ACKNOWLEDGEMENTS

The authors would like to thank current and former Nair lab members, Dr. Zachary J. S. Mays, Dr. Debika Choudhury, Sean F. Sullivan, Trevor B. Nicks, Rana Said, Maya Vaishnaw, and Alexis Barselau for helpful discussions.

## AUTHOR CONTRIBUTIONS

N.U.N., K.M., T.C.C., and V.D.T. conceived the idea, N.U.N., T.C.C., V.D.T. designed the research project and T.C.C., V.D.T., N.B.K., A.S., G.G.S., P.K.R., N.U.N. co-wrote the manuscript. T.C.C., V.D.T., K.M., and A.R. performed the experiments. T.C.C. and V.D.T. analyzed the data. N.B.K., A.S., G.G.S., P.K.R. performed analysis of the computational results. All the authors have reviewed the manuscript and approved it for submission.

## REFERENCES

1 Kong, J.-Q. Phenylalanine ammonia-lyase, a key component used for phenylpropanoids production by metabolic engineering. RSC advances 5, 62587–62603 (2015).

2 Parmeggiani, F., Weise, N. J., Ahmed, S. T. & Turner, N. J. Synthetic and therapeutic applications of ammonia-lyases and aminomutases. Chemical reviews 118, 73–118 (2018).

3 Klumbys, E., Zebec, Z., Weise, N. J., Turner, N. J. & Scrutton, N. S. Bio-derived production of cinnamyl alcohol via a three step biocatalytic cascade and metabolic engineering. Green Chemistry 20, 658–663 (2018).

4 Toogood, H. S. & Scrutton, N. S. Discovery, characterization, engineering, and applications of ene-reductases for industrial biocatalysis. ACS catalysis 8, 3532–3549 (2018).

5 Parmeggiani, F., Lovelock, S. L., Weise, N. J., Ahmed, S. T. & Turner, N. J. Synthesis of D-and l-phenylalanine derivatives by phenylalanine ammonia lyases: a multienzymatic cascade process. Angewandte Chemie 127, 4691–4694 (2015).

6 Weise, N. J. et al. Zymophore identification enables the discovery of novel phenylalanine ammonia lyase enzymes. Scientific reports 7, 1–9 (2017).

7 Zhang, F., Ren, J. & Zhan, J. Identification and Characterization of an Efficient Phenylalanine Ammonia-Lyase from Photorhabdus luminescens. Applied Biochemistry and Biotechnology 193, 1099–1115 (2021).

8 Moffitt, M. C. et al. Discovery of two cyanobacterial phenylalanine ammonia lyases: kinetic and structural characterization. Biochemistry 46, 1004–1012 (2007).

9 Isabella, V. M. et al. Development of a synthetic live bacterial therapeutic for the human metabolic disease phenylketonuria. Nature biotechnology 36, 857–864 (2018).

10 Burton, B. K. et al. Pegvaliase for the treatment of phenylketonuria: Results of the phase 2 dose-finding studies with long-term follow-up. Molecular genetics and metabolism 130, 239–246 (2020).

11 Yang, J. et al. Thermosensitive micelles encapsulating phenylalanine ammonia lyase act as a sustained and efficacious therapy against colorectal cancer. Journal of biomedical nanotechnology 15, 717–727 (2019).

12 Babich, O. O., Pokrovsky, V. S., Anisimova, N. Y., Sokolov, N. N. & Prosekov, A. Y. Recombinant l-phenylalanine ammonia lyase from Rhodosporidium toruloides as a potential anticancer agent. Biotechnology and Applied Biochemistry 60, 316–322 (2013).

13 Bartsch, S. & Bornscheuer, U. T. Mutational analysis of phenylalanine ammonia lyase to improve reactions rates for various substrates. Protein Engineering, Design & Selection 23, 929–933 (2010).

14 Bencze, L. C. et al. Expanding the substrate scope of phenylalanine ammonia-lyase from Petroselinum crispum towards styrylalanines. Organic & biomolecular chemistry 15, 3717–3727 (2017).

15 Nagy, E. Z. et al. Mapping the hydrophobic substrate binding site of phenylalanine ammonia-lyase from Petroselinum crispum. ACS Catalysis 9, 8825–8834 (2019).

16 Flachbart, L. K., Sokolowsky, S. & Marienhagen, J. Displaced by deceivers: prevention of biosensor cross-talk is pivotal for successful biosensor-based high-throughput screening campaigns. ACS synthetic biology 8, 1847–1857 (2019).

17 Mays, Z. J., Mohan, K., Trivedi, V. D., Chappell, T. C. & Nair, N. U. Directed evolution of Anabaena variabilis phenylalanine ammonia-lyase (PAL) identifies mutants with enhanced activities. Chemical Communications 56, 5255–5258 (2020).

18 Fowler, D. M. et al. High-resolution mapping of protein sequence-function relationships. Nature methods 7, 741 (2010).

19 Hietpas, R. T., Jensen, J. D. & Bolon, D. N. Experimental illumination of a fitness landscape. Proceedings of the National Academy of Sciences 108, 7896–7901 (2011).

20 Wrenbeck, E. E., Faber, M. S. & Whitehead, T. A. Deep sequencing methods for protein engineering and design. Current opinion in structural biology 45, 36–44 (2017).

21 Wrenbeck, E. E., Azouz, L. R. & Whitehead, T. A. Single-mutation fitness landscapes for an enzyme on multiple substrates reveal specificity is globally encoded. Nature communications 8, 1–10 (2017).

22 Jones, E. M. et al. Structural and functional characterization of G protein–coupled receptors with deep mutational scanning. Elife 9, e54895 (2020).

23 McLaughlin Jr, R. N., Poelwijk, F. J., Raman, A., Gosal, W. S. & Ranganathan, R. The spatial architecture of protein function and adaptation. Nature 491, 138–142 (2012).

24 Araya, C. L. et al. A fundamental protein property, thermodynamic stability, revealed solely from large-scale measurements of protein function. Proceedings of the National Academy of Sciences 109, 16858–16863 (2012).

25 Bata, Z. et al. Substrate Tunnel Engineering Aided by X-ray Crystallography and Functional Dynamics Swaps the Function of MIO-Enzymes. ACS Catalysis 11, 4538–4549 (2021).

26 Brenan, L. et al. Phenotypic characterization of a comprehensive set of MAPK1/ERK2 missense mutants. Cell reports 17, 1171–1183 (2016).

27 Calabrese, J. C., Jordan, D. B., Boodhoo, A., Sariaslani, S. & Vannelli, T. Crystal structure of phenylalanine ammonia lyase: multiple helix dipoles implicated in catalysis. Biochemistry 43, 11403–11416 (2004).

28 Heberling, M. M. et al. Ironing out their differences: dissecting the structural determinants of a phenylalanine aminomutase and ammonia lyase. ACS chemical biology 10, 989–997 (2015).

29 Hermes, J. D., Weiss, P. M. & Cleland, W. Use of nitrogen-15 and deuterium isotope effects to determine the chemical mechanism of phenylalanine ammonia-lyase. Biochemistry 24, 2959–2967 (1985).

30 Jun, S.-Y. et al. Biochemical and structural analysis of substrate specificity of a phenylalanine ammonia-lyase. Plant physiology 176, 1452–1468 (2018).

31 Louie, G. V. et al. Structural determinants and modulation of substrate specificity in phenylalanine-tyrosine ammonia-lyases. Chemistry & biology 13, 1327–1338 (2006).

32 Melnikov, A., Rogov, P., Wang, L., Gnirke, A. & Mikkelsen, T. S. Comprehensive mutational scanning of a kinase in vivo reveals substrate-dependent fitness landscapes. Nucleic acids research 42, e112–e112 (2014).

33 Ritter, H. & Schulz, G. E. Structural basis for the entrance into the phenylpropanoid metabolism catalyzed by phenylalanine ammonia-lyase. The Plant Cell 16, 3426–3436 (2004).

34 Rockah-Shmuel, L., Tóth-Petróczy, Á. & Tawfik, D. S. Systematic mapping of protein mutational space by prolonged drift reveals the deleterious effects of seemingly neutral mutations. PLoS Comput Biol 11, e1004421 (2015).

35 Starita, L. M. et al. Activity-enhancing mutations in an E3 ubiquitin ligase identified by high-throughput mutagenesis. Proceedings of the National Academy of Sciences 110, E1263–E1272 (2013).

36 Wang, L. et al. Structural and biochemical characterization of the therapeutic Anabaena variabilis phenylalanine ammonia lyase. Journal of molecular biology 380, 623–635 (2008).

37 Wang, L. et al. Structure-based chemical modification strategy for enzyme replacement treatment of phenylketonuria. Molecular genetics and metabolism 86, 134–140 (2005).

38 Cooke, H. A., Christianson, C. V. & Bruner, S. D. Structure and chemistry of 4-methylideneimidazole-5-one containing enzymes. Current opinion in chemical biology 13, 460–468 (2009).

39 Feng, L., Wanninayake, U., Strom, S., Geiger, J. & Walker, K. D. Mechanistic, mutational, and structural evaluation of a Taxus phenylalanine aminomutase. Biochemistry 50, 2919–2930 (2011).

40 Cooke, H. A. & Bruner, S. D. Probing the active site of MIO-dependent aminomutases, key catalysts in the biosynthesis of β-amino acids incorporated in secondary metabolites. Biopolymers 93, 802–810 (2010).

41 Tomoiagă, R. B. et al. Saturation Mutagenesis for Phenylalanine Ammonia Lyases of Enhanced Catalytic Properties. Biomolecules 10, 838 (2020).

42 MacDonald, M. J. & D’Cunha, G. B. A modern view of phenylalanine ammonia lyase. Biochemistry and Cell Biology 85, 273–282 (2007).

43 Sato, T., Kiuchi, F. & Sankawa, U. Inhibition of phenylalanine ammonia-lyase by cinnamic acid derivatives and related compounds. Phytochemistry 21, 845–850 (1982).

44 Zoń, J. & Laber, B. Novel phenylalanine analogues as putative inhibitors of enzymes acting on phenylalanine. Phytochemistry 27, 711–714 (1988).

45 Poppe, L. & Rétey, J. Friedel–crafts-type mechanism for the enzymatic elimination of ammonia from histidine and phenylalanine. Angewandte Chemie International Edition 44, 3668–3688 (2005).

46 Phillips, J. C. et al. Scalable molecular dynamics on CPU and GPU architectures with NAMD. The Journal of chemical physics 153, 044130 (2020).

47 Pinto, G. P. et al. New insights in the catalytic mechanism of tyrosine ammonia-lyase given by QM/MM and QM cluster models. Archives of biochemistry and biophysics 582, 107–115 (2015).

48 Bushnell, B., Rood, J. & Singer, E. BBMerge–Accurate paired shotgun read merging via overlap. PloS one 12, e0185056 (2017).

49 Langmead, B. & Salzberg, S. L. Fast gapped-read alignment with Bowtie 2. Nature methods 9, 357 (2012).

50 Webb, B. & Sali, A. Comparative protein structure modeling using MODELLER. Current protocols in bioinformatics 54, 5.6. 1–5.6. 37 (2016).

51 Morris, G. M. et al. AutoDock4 and AutoDockTools4: Automated docking with selective receptor flexibility. Journal of computational chemistry 30, 2785–2791 (2009).

52 Lindorff-Larsen, K. et al. Improved side-chain torsion potentials for the Amber ff99SB protein force field. Proteins: Structure, Function, and Bioinformatics 78, 1950–1958 (2010).

53 Berendsen, H. J., van der Spoel, D. & van Drunen, R. GROMACS: a message-passing parallel molecular dynamics implementation. Computer physics communications 91, 43–56 (1995).

54 Van Der Spoel, D. et al. GROMACS: fast, flexible, and free. Journal of computational chemistry 26, 1701–1718 (2005).

55 Pronk, S. et al. GROMACS 4.5: a high-throughput and highly parallel open source molecular simulation toolkit. Bioinformatics 29, 845–854 (2013).

56 Páll, S., Abraham, M. J., Kutzner, C., Hess, B. & Lindahl, E. in International conference on exascale applications and software. 3–27 (Springer).

57 Abraham, M. J. et al. GROMACS: High performance molecular simulations through multi-level parallelism from laptops to supercomputers. SoftwareX 1, 19–25 (2015).

58 Berendsen, H. J., Postma, J. v., van Gunsteren, W. F., DiNola, A. & Haak, J. R. Molecular dynamics with coupling to an external bath. The Journal of chemical physics 81, 3684–3690 (1984).

59 Darden, T., York, D. & Pedersen, L. Particle mesh Ewald: An N· log (N) method for Ewald sums in large systems. The Journal of chemical physics 98, 10089–10092 (1993).

60 Melo, M. C. et al. NAMD goes quantum: an integrative suite for hybrid simulations. Nature methods 15, 351 (2018).

61 MOPAC2016 (Colorado Springs, CO, USA, 2016).

62 Neese, F. Wiley Interdiscip. Rev.: Comput. Mol. Sci. 2, 73–78. (2018).

63 Fiorin, G., Klein, M. L. & Hénin, J. Using collective variables to drive molecular dynamics simulations. Molecular Physics 111, 3345–3362 (2013).

