## Supplemental Information for "In-depth sequence-function characterization reveals multiple paths to enhance phenylalanine ammonia-lyase (PAL) activity"

Vikas D. Trivedi[1]*, Todd C. Chappell[1]*, Naveen B. Krishna[2], Anuj Shetty[2], Gladstone G. Sigamani[2], Karishma Mohan[1], Athreya Ramesh[1], Pravin Kumar R.[2], Nikhil U. Nair[1✉]
[1] Department of Chemical and Biological, Tufts University, Medford, USA 02155
[2] Kcat Enzymatic Private Limited, Bengaluru, Karnataka, India
*Equal contribution

#### SUPPLEMENTARY FIGURES.

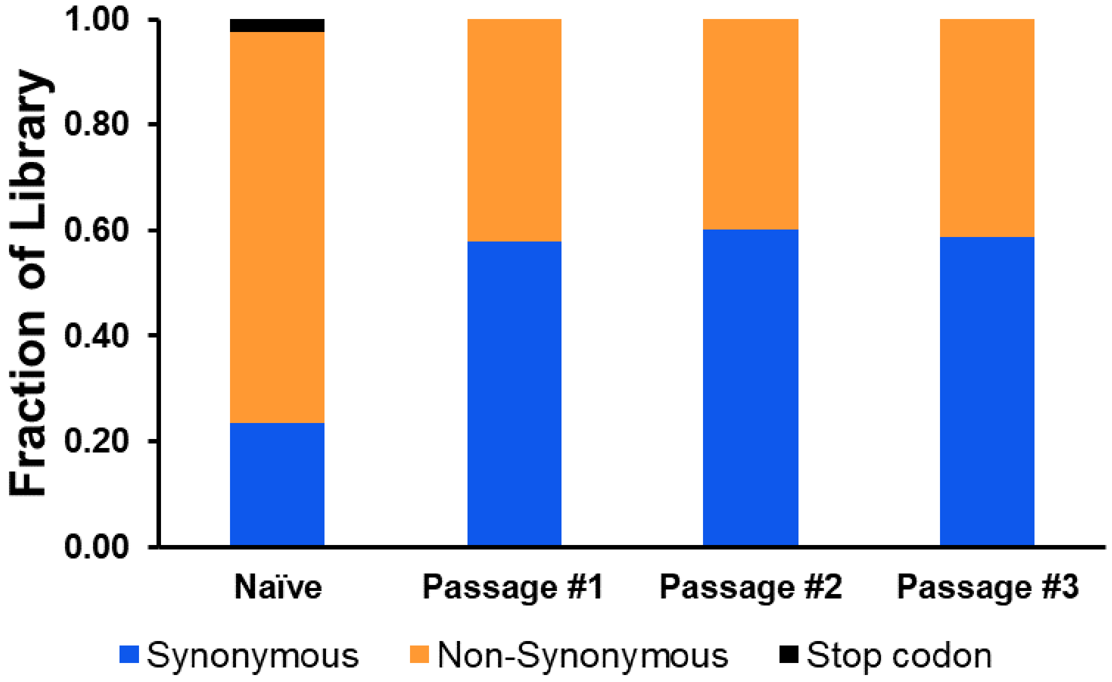

**Figure S1. Types of mutations in passages.** Mutation type measured across enrichment passages. Stop codons are black, synonymous mutations are blue, and non-synonymous mutations are orange.

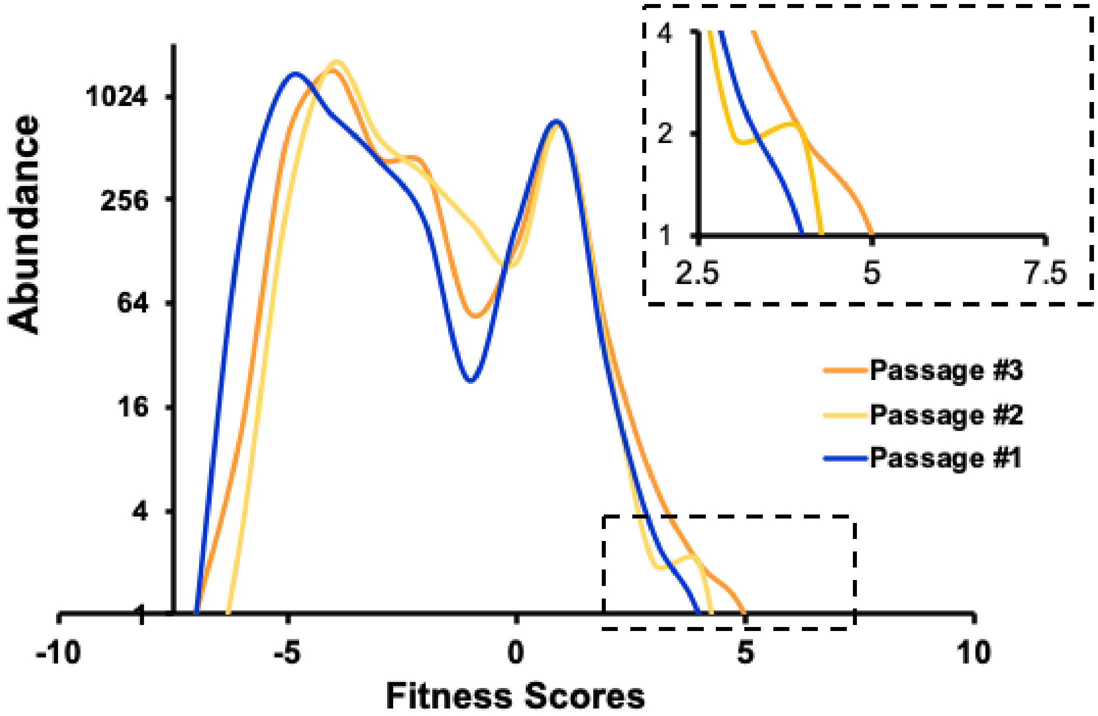

**Figure S2. Fitness distribution during passages.** Fitness for all three passages was calculated relative to the Naïve population. Inset shows blowup of highly fit mutations. Passage #1 is blue, Passage #2 is yellow, and Passage #3 is orange.

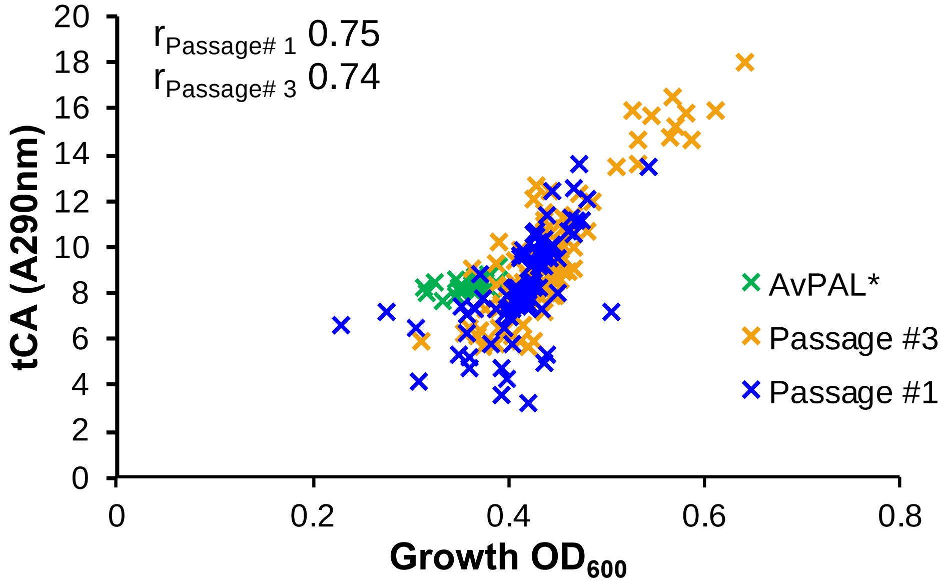

**Figure S3. Growth and tCA production during passages.** Growth represented as OD600 and tCA production after 24 h for individual colonies randomly picked from Passage #1 (blue), Passage #3 (orange) and AvPAL* (green). Spearman correlation coefficients for data points from Passage#1 and #3 are indicated in the inset.

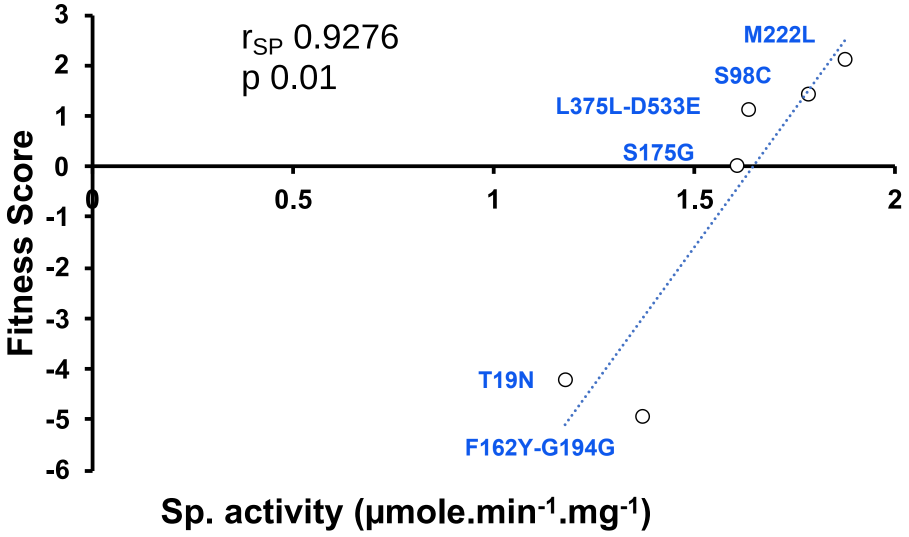

**Fig. S4. Correlation between specific activity and fitness score.** The specific activity of the single mutants characterized in Mays et al., 2020 were correlated with the fitness sores of the Ep-PCR library after enrichment. The data supports use of fitness as proxy for enzyme activity. Spearman correlation coefficient is indicated in the inset along with p-value.

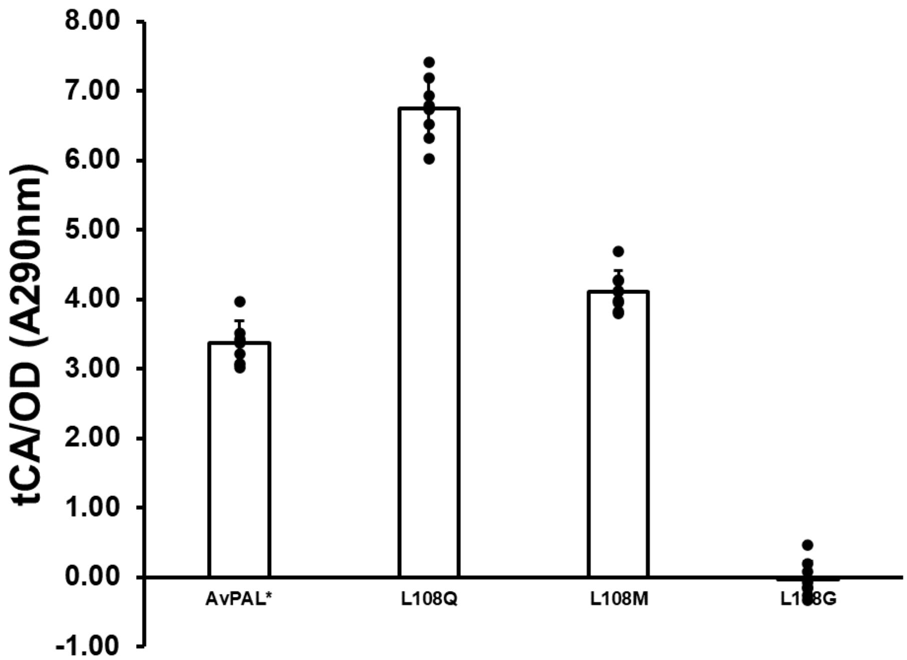

**Figure S5. Whole cell tCA measurement of AvPAL*, L108Q, M, and G.** The data supports DMS fitness data and use of L108G as negative control enzyme. One standard deviation is indicated as the error bars, n = 6.

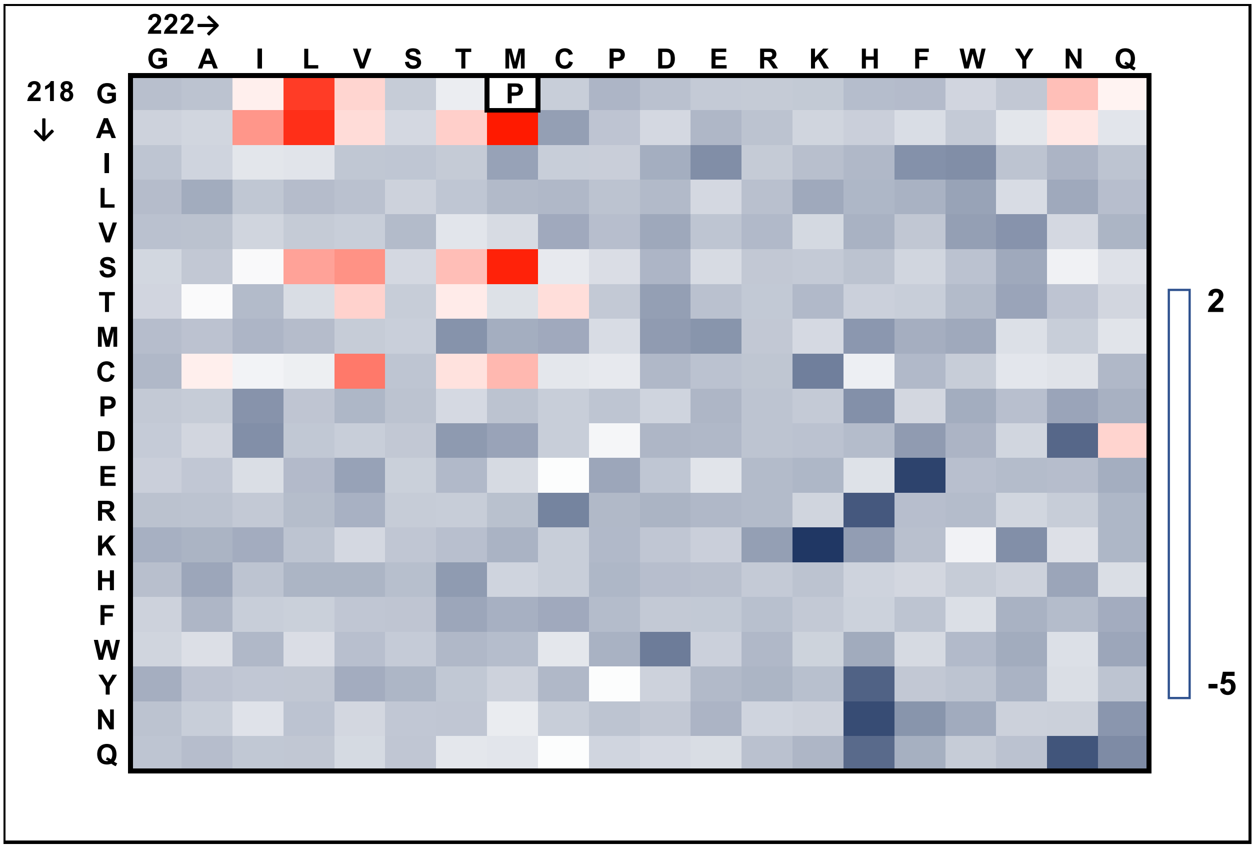

**Figure S6. Fitness map of AvPAL* 218X-222X double mutant library**. Data generated after enrichment on phenylalanine. The data supports that G218 and M222 mutations are largely epistatic and co-mutation of both does not yield variants with activity higher than single mutants. P = parental AvPAL* sequence.

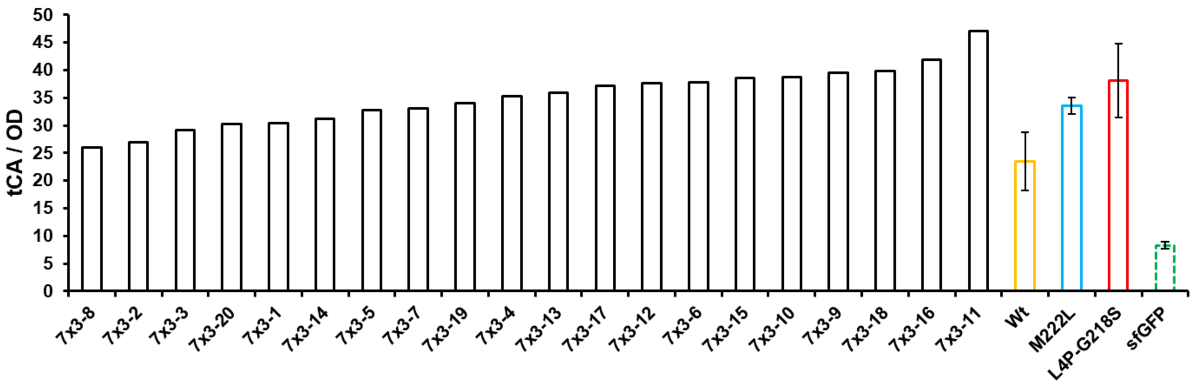

**Figure S7. tCA/OD measurement of randomly picked colonies from $^{7}C_{3}$ library after enrichment on Phe.** All variants were active, demonstrating at least parental level of tCA production. Control mutants chosen for comparison since they are the most active variants isolated previously (Mays, Mohan, et al, 2020). One standard deviation is indicated as the error bars, n = 3.

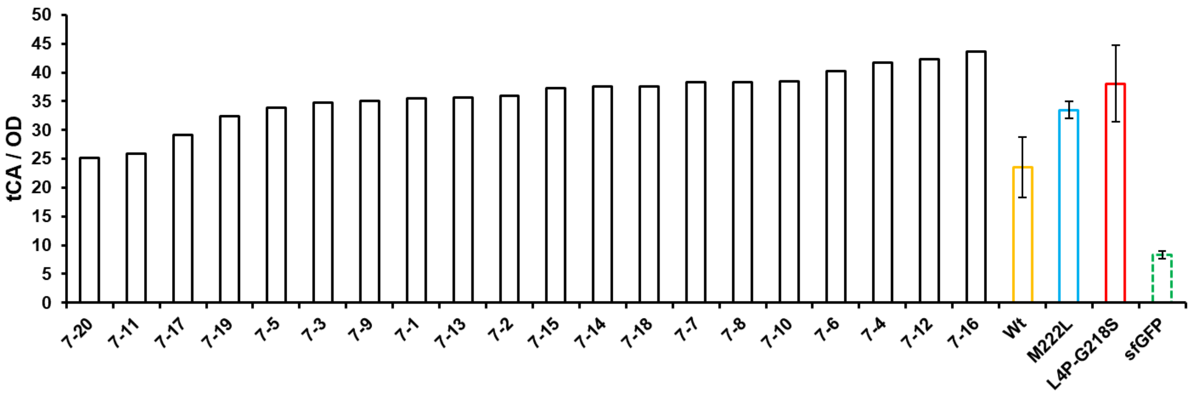

**Figure S8. tCA/OD measurement of randomly picked colonies from $^{7}C_{7}$ library after enrichment on Phe.** All variants were active, demonstrating at least parental level of tCA production. Control mutants chosen for comparison since they are the most active variants isolated previously (Mays, Mohan, et al, 2020). One standard deviation is indicated as the error bars, n = 3.

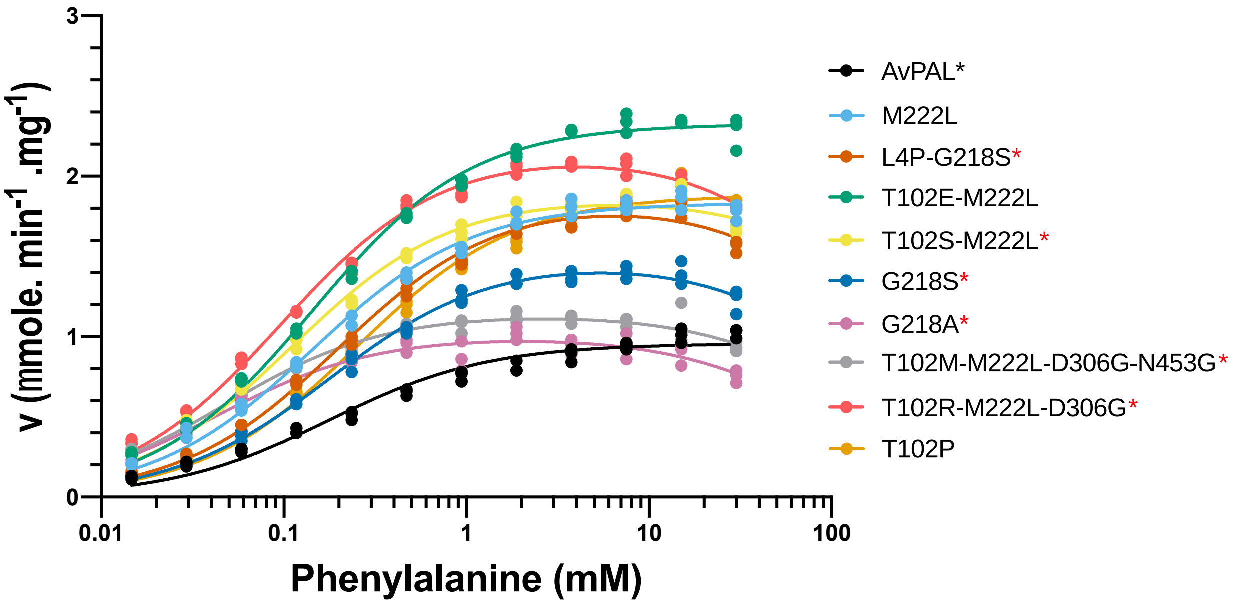

**Figure S9. Michaelis-Menten plots for AvPAL* variants.** All mutants have higher activity than parental at physiologically relevant Phe concentrations (<1 mM). L4P-G218, and M222L were previously isolated (Mays, Mohan, et al, 2020). * indicates substrate inhibition. One standard deviation is indicated as the error bars, n = 3.

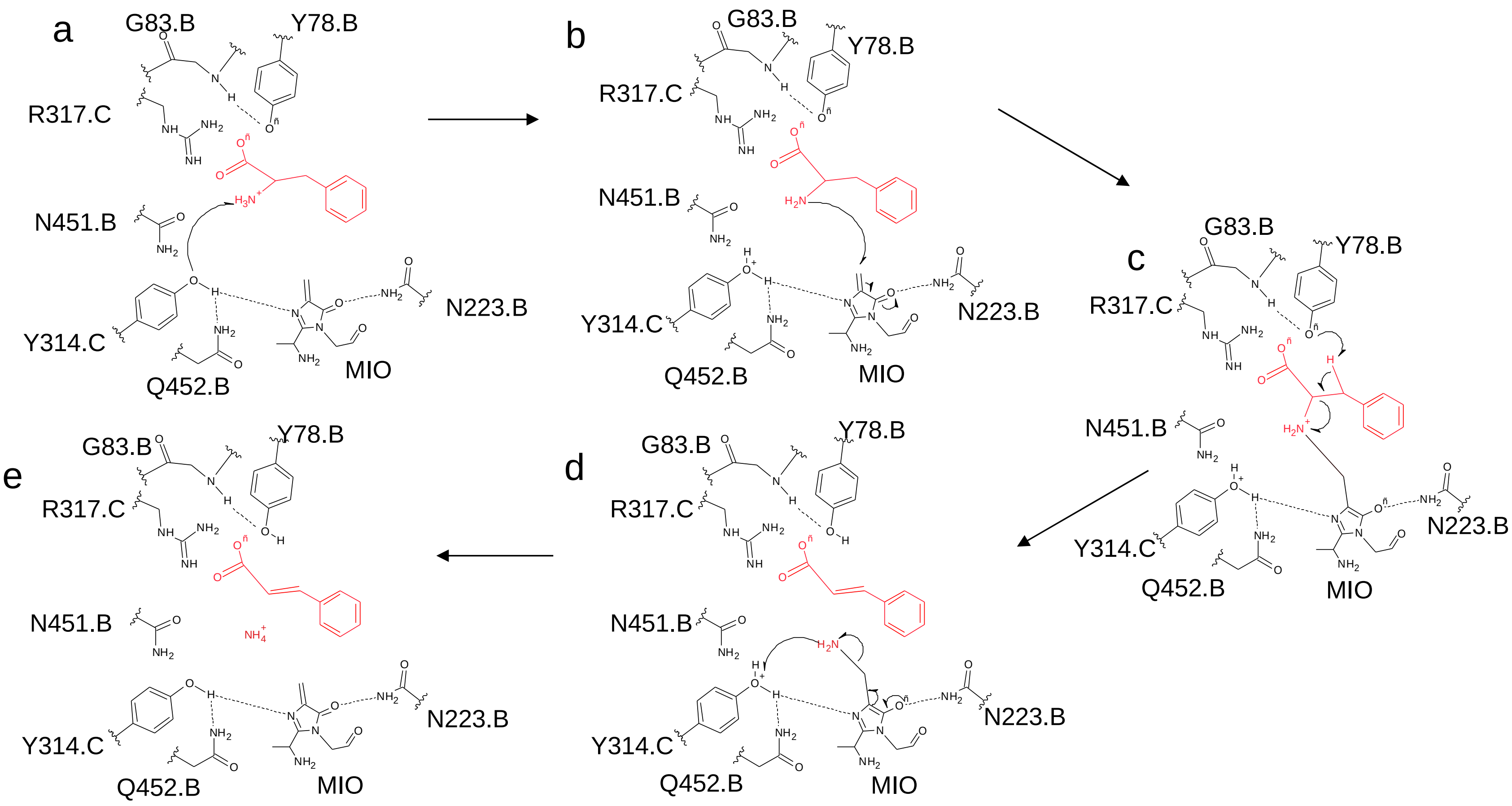

**Figure S10. Schematic representation of PAL mechanism** (adapted from Jun et al 2018). **a−e)** N-MIO reaction pathway from substrate to product formation. The substrate, Phe, is represented in red and the catalytic residues of the active site are in black.

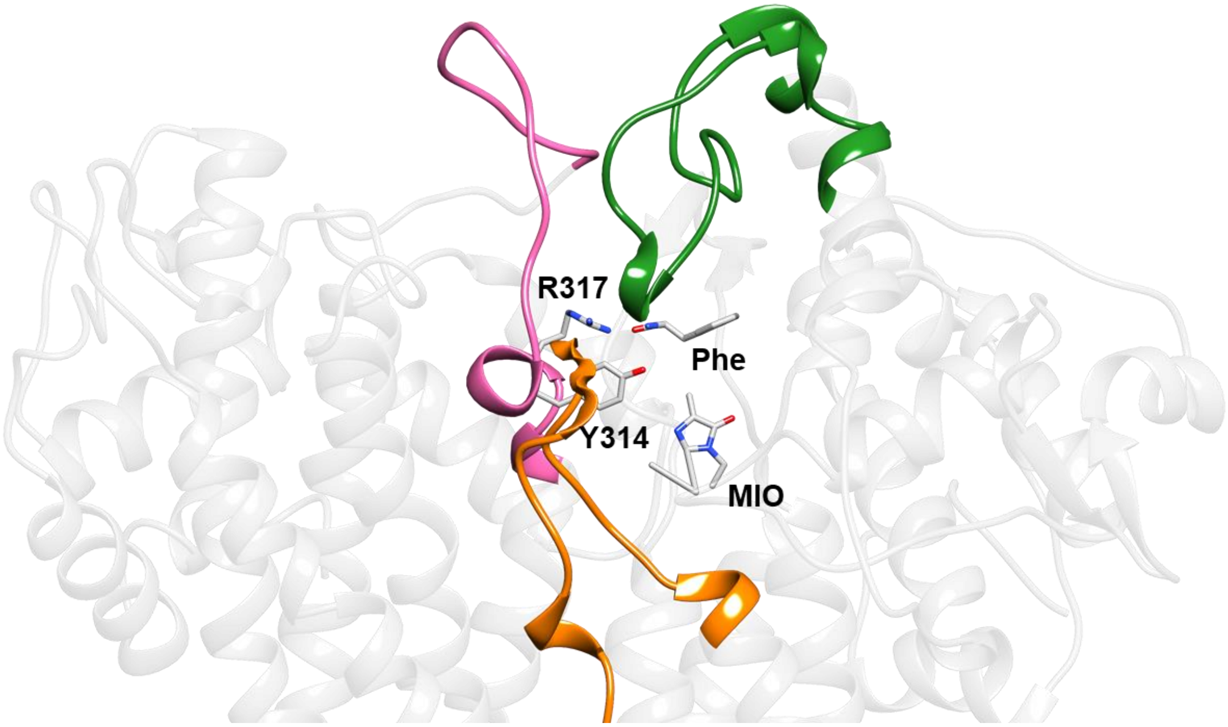

**Figure S11. Regions of the enzyme showing largest RMSF.** The regions of major fluctuation in green, magenta and orange mapped over the 3D structure. The regions depicted as coloured ribbons are derived from the regions enclosed in boxes in the RMSF graph in (**Fig 5g**).

**Metadynamics simulation predict the free energy landscapes in the active site of protein.**

In metadynamics based approaches, choice of a collective variable (CVs) in the design of experiment is crucial. Using three CVs which includes the distance between the center of mass (COM) of substrate atoms and COM of Y314 (CV1), COM of heavy atoms in the backbone of residues in conserved secondary structures that were present within 5 Å of G218, M222 and L108 residues (CV2) (**Fig. S12a**). The choice of picking the region around the mutations to define a CV is novel. The intramolecular network formed by the residues in this region is expected to change due to mutations and consequently, affect the substrate binding dynamics. The Free Energy Surface (FES) obtained from different sets of experiment provide us an understanding of the energy bins associated within the active site and the regions around it. The closer the minima are towards the origin (lower-left corner), better is the stability of the system, which is seen for the hyperactive variants but not the L108G negative control (**Fig. S12b-g**).

The conformational changes of the substrate in the minima were traced back and compared between AvPAL* and mutants (**Fig. S12h-m**). One interaction that we repeatedly observe in the mutants is that of substrate with Y78, a crucial residue that desaturates the substrate. Overall, Phe was better stabilized by H-bond (green dotted lines), alkyl-π (pink dotted lines) and electrostatic interactions (orange dotted lines) in the hyperactive mutants than by parental AvPAL*. The conformations of substrate in the mutant active site superimposed over the AvPAL* conformation is shown in **Fig. S12n**.

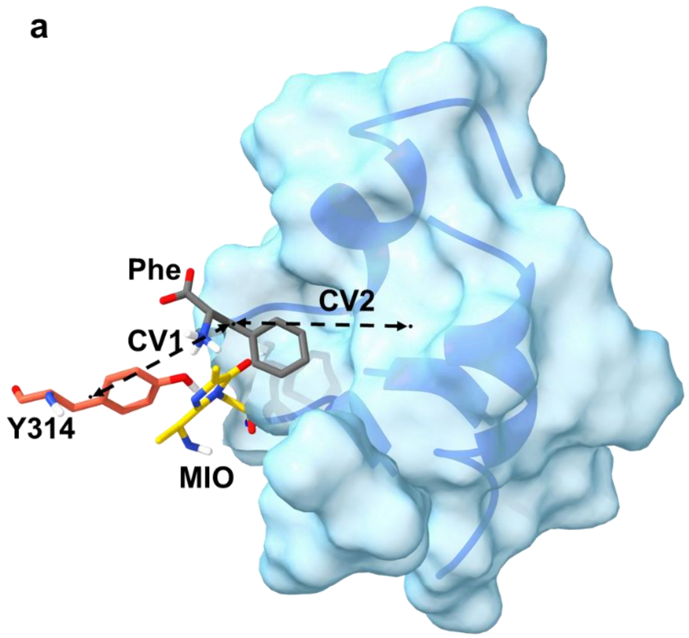

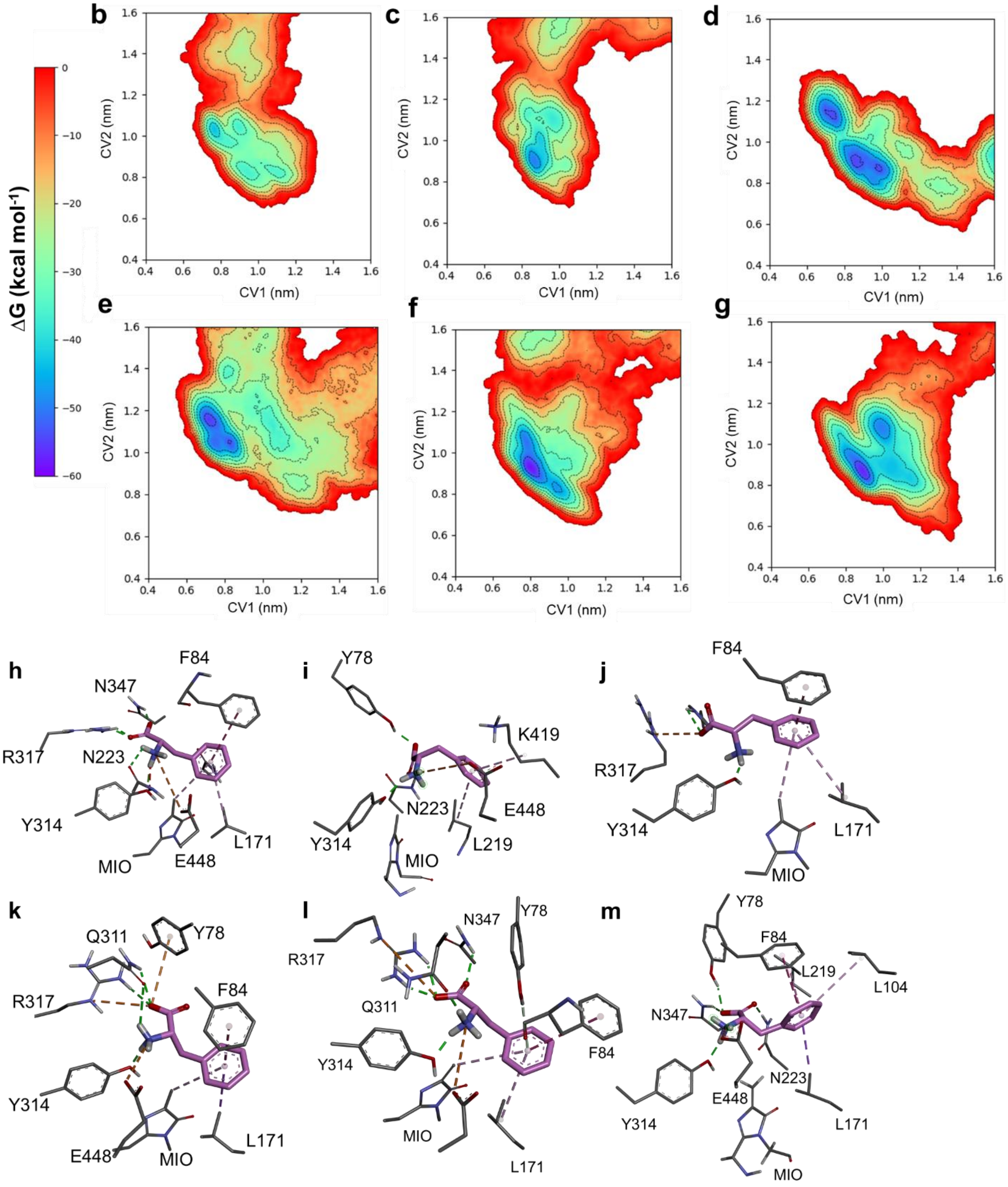

ΔG (kcal mol-1)
0
−10
−20
−30
−40
−50
−60
b
c
d
e
f
g
CV1 (nm)
CV2 (nm)
h
F84
N347
R317
N223
Y314
MIO
E448
L171
i
Y78
K419
N223
E448
Y314
MIO
L219
j
F84
R317
Y314
L171
MIO
k
Q311
Y78
R317
F84
Y314
E448
MIO
L171
l
N347
Y78
R317
Q311
Y314
F84
MIO
L171
m
Y78
F84
L219
L104
N347
N223
E448
Y314
L171
MIO

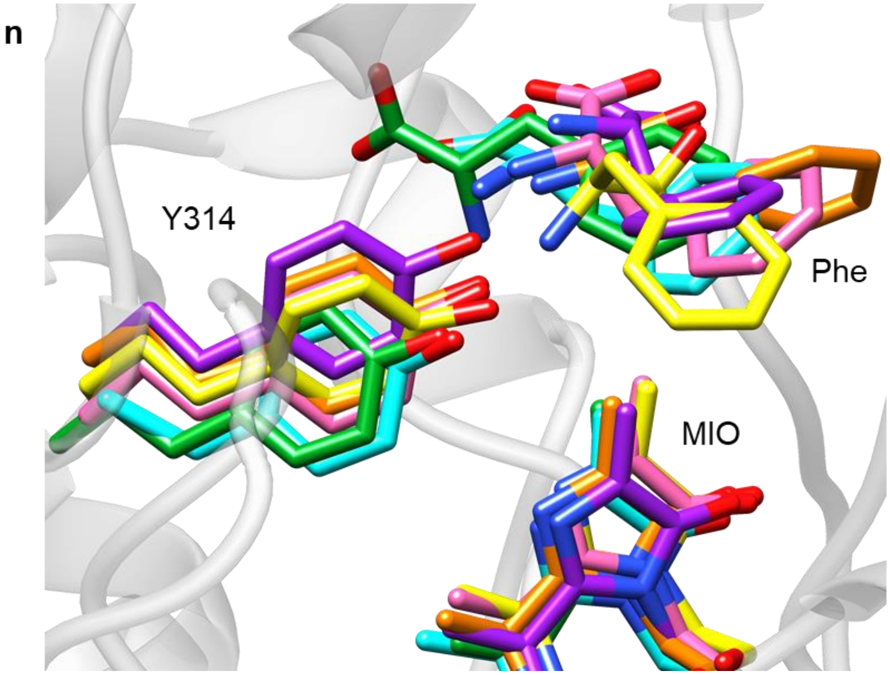

**Figure S12. Results from MD metadynamic studies. a)** Schematic representation of the two CVs (CV1 and CV2) that were used in the metadynamics simulation. Substrate is shown in grey sticks and MIO cofactor in yellow stick model. The region 5 Å around the mutation sites is represented as blue surface. The FES for **b)** AvPAL*, **c)** G218S, **d)** M222L, **e)** L108G, **f)** T102E-M222L, and **g)** T102R-M222L-D306G, respectively. FES profile was computed for CVs CV1 and CV2. CV1 is the distance between COM of Substrate and COM of Y314 in x−axis. CV2 is the distance between COM derived from backbone atoms of the residue within 5Å of residues M222, G218, L108 and COM of substrate in y-axis. AvPAL* and L108G showed minima away from the plausible attack conformation whereas the other mutants showed lesser minima and much closer to Y314. Intermolecular interactions between the enzyme and the substrate derived from the minima of the respective simulation experiments shown in (b-g) for **h)** AvPAL*, **i)** G218S, **j)** M222L, **k)** L108G, **l)** T102E-M222L, and **m)** T102R-M222L-D306G. **n)** Superimposition of the FES minima of parental and the mutant E-S complexes. Elementary stick model coloured as cyan, yellow, pink, green, purple, orange represents the minima of the substrate in AvPAL*, G218S, M222L, L108G, T102E-M222L and T102R-M222L-D306G, respectively.

**Steered molecular dynamics (SMD) studies show steady and seamless diffusion of the substrate in the mutant, N453S.**

The unbinding process of the substrate in wild and N453S shows rupture force of 600 kJ $mol^{-1}$ $nm^{-1}$ and 400 kJ $mol^{-1}$ $nm^{-1}$ and dissociation at 4.2 ns and 3 ns, respectively (**Fig. S13a, c**). This signifies a faster dissociation of the substrate in N453S. The reassociation of the substrate in wild and N453S simulations conducted with negative force, where -30 kJ $mol^{-1}$ $nm^{-1}$ was used as the primary force constant. The reassociation of the substrate happened faster in N453S than in AvPAL*, which is 4.5 ns and 6.5 ns, respectively. During the reassociation process, we observed that the substrate approached ~4 Å from the active site for in N453S within 4.5 ns, which took the parental enzyme 8.5 ns. Diffusion snapshots of Phe exiting and entering the active site of the AvPAL* are visualized in **Movie S1, S2**.

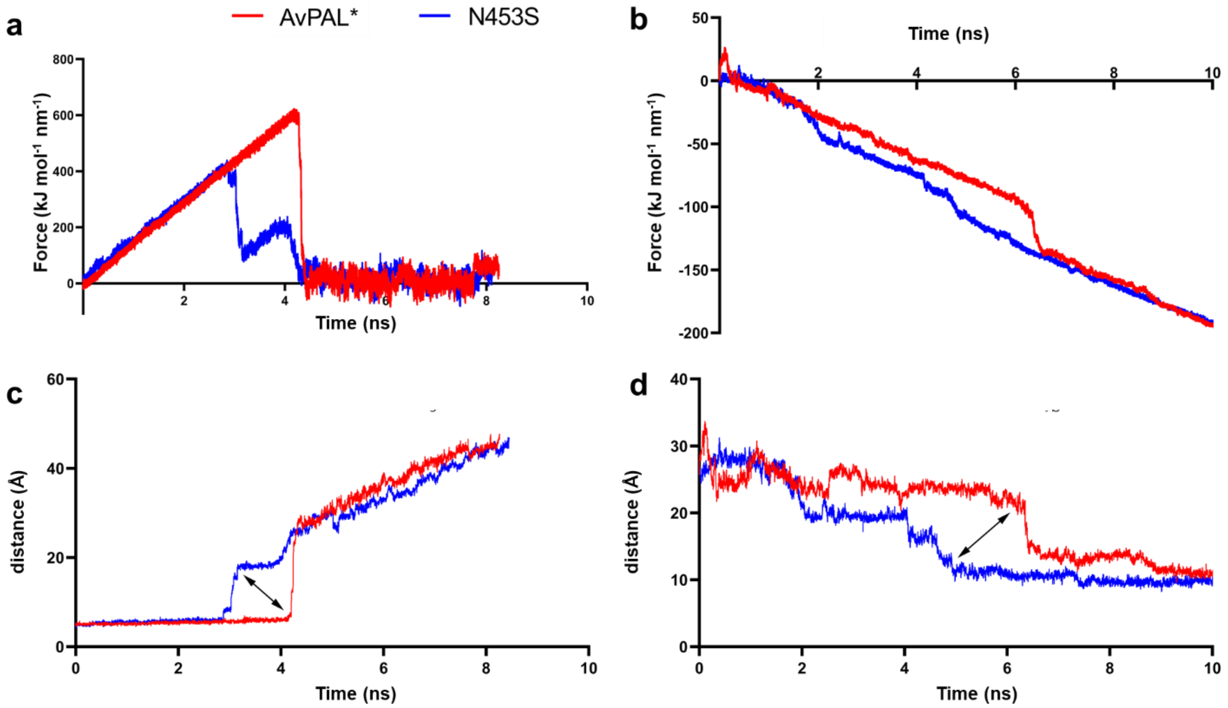

**Figure S13. SMD profile for substrate unbinding and reassociation. a, b)** The time-dependent external force applied. **c, d)** Distance calculated between MIO(Cβ2) and Phe(N) atoms during unbinding and re-association/entry process, respectively.

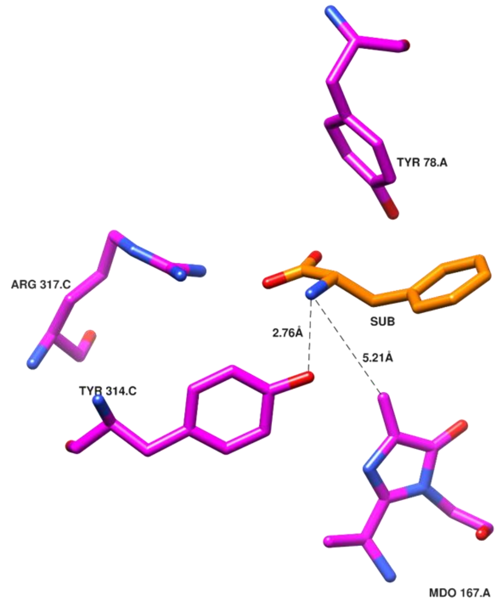

**Figure S14. Schematic representation of Michaelis complex of PAL.** Substrate Phe highlighted in orange and active site residues in magenta.

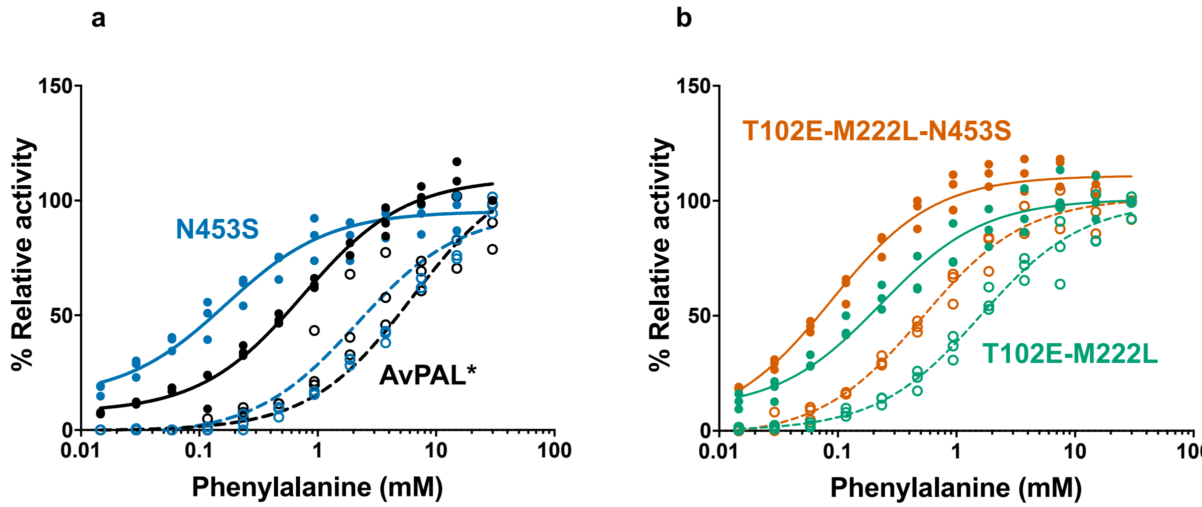

**Figure S15**. **Effect of N453S on product inhibition.** Activity of **a)** AvPAL*, N453S and **b)** T102E-M222L, T102E-M222L-N453S on varying concentration Phe (0−30 mM) in presence (empty circles and dashed line) and absence (filled circles and solid line) of tCA (150 µM). In general, N453S variants showed lower tCA inhibition compared to their parental counterparts.

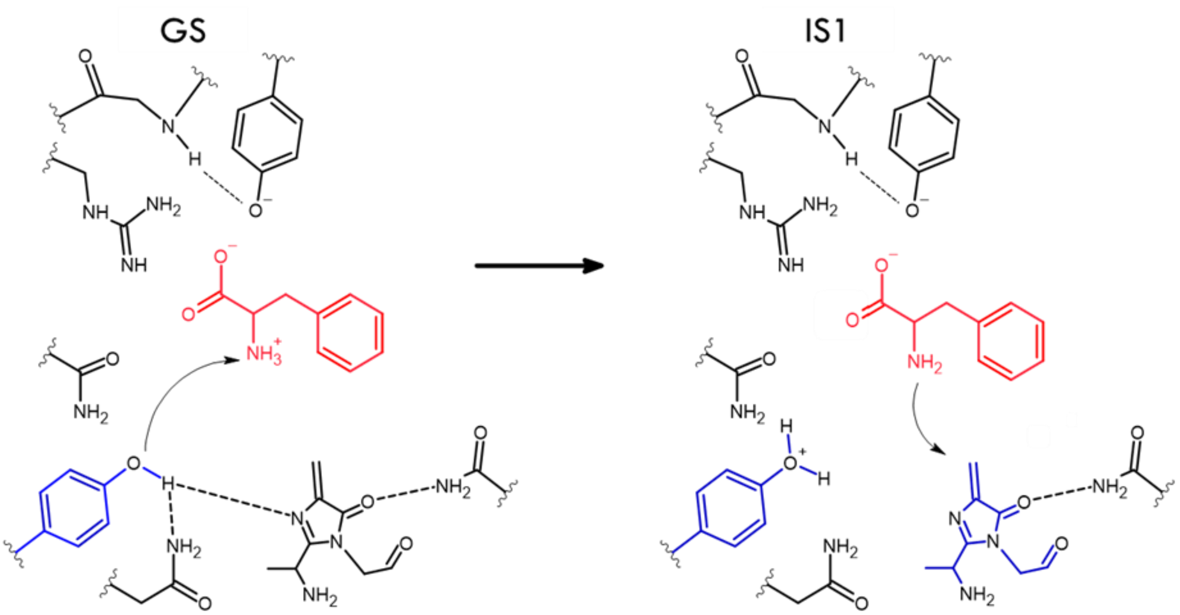

**Figure S16. First step of N-MIO reaction pathway.** As Phe enters the active site and near attack conformation is attained, the carboxyl group forms charged electrostatic interactions with R317 of the adjacent chain. Y314 of the adjacent chain abstracts a proton from the amino group of the substrate depicted in GS followed by the formation of attack conformations formed between the substrate and MIO shown in IS1[1].

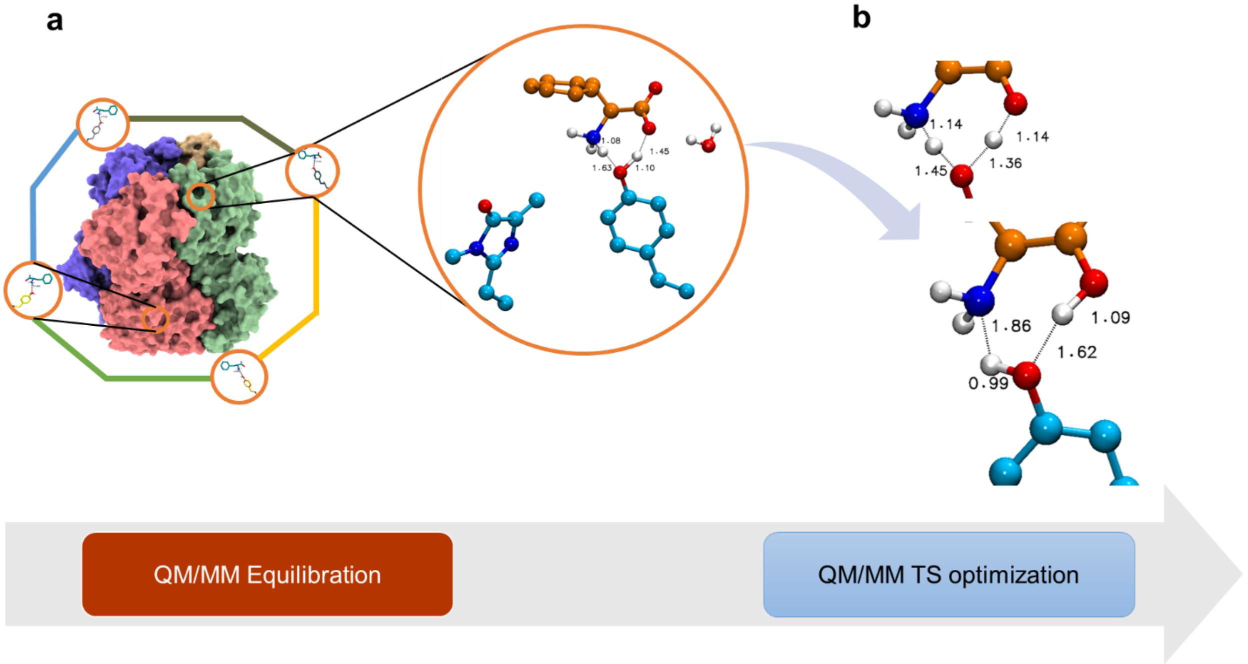

**Figure S17. QM/MM implemented in this study. a)** 1 µs MD equilibrated E-S conformation was extended with QMMM-PM7 simulation using NAMD 2.13 to derive the attack conformation. This reduced the distance between Tyr hydroxyl and Phe amine from ~3.5 Å to 1.5 Å in all the 4 chains. **b)** Post this, DFT level TS optimizations were adapted to obtain transition state and the final state.

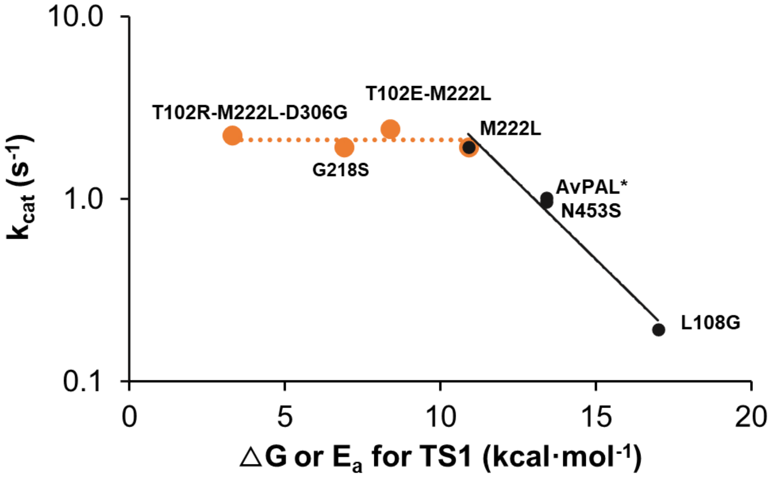

**Figure S18. Plot of experimentally determined $k_{cat}$ versus free/activation energy (ΔG or $E_a$) of transition state 1 (TS1).** The Arrhenius equation ($k = A \cdot \exp(-E_a/RT)$) relates reaction rate to the activation energy ($E_a$) of the rate-limiting step. There is no significant increase in reaction rate beyond ~2 $s^{-1}$, indicating formation of intermediate state 1 (IS1) through TS1 is no longer rate limiting.

**Changes observed in QM/MM transition states across mutants.**

After the formation of the first intermediate state (IS1) for the first step of the reaction (proton abstraction from Y314) the system undergoes certain atomistic changes to stabilize the new-formed states. The changes between AvPAL*, L108G and T102R-M222L-D306G were considered to make observations. T102R-M222L-D306G is used as representative for the other hyperactive mutants. Noticeable changes around the active site residues during this IS1 state can be observed in residues L104, Y78 & M416 **(Fig. S19a)**. In the case of Y78, the side chain is in a different orientation when compared to the rest of the mutants. M416 is closer to the Phe in the hyperactive variants when compared to the AvPAL* and L108G. L104 is also pushed slightly closer to the Phe in the L108G mutant. These minor changes in residue interactions leads to a change in the Phe conformation during the IS1 state. The Phe amino group is placed at 3.96 Å, 3.62 Å & 4.66 Å from the methylidene carbon of the MIO cofactor in AvPAL*, Hyperactive variants and L108G, respectively. Distances for the hyperactive variants were around 3.65 Å in all cases. After the first step of the reaction, in the negative variant (L108G) the substrate takes a non-catalytic conformation that prevents it from moving closer to MIO and stays 1 Å away for a longer period and moves further away from MIO and stays in that conformation throughout. The difference in the rotation of the bond calculated between 4 atoms yields -66.18°,152.11° and -91.10° for the parental, hyperactive, and negative control mutants, respectively **(Fig. S19b)**. The difference in the bond rotation is sufficient for the L108G mutant to yield a low activity conformation. L108G mutation changes the hydrophobic nature around the residue L104, focusing its interaction towards Phe when compared to the rest. As seen from the space filling model in **Fig. S19c**, L104 occupies more space towards the head region of the Phe thereby adding to the change in conformation in the L108G variant. The stability of phenyl moiety of the substrate in the active site is impacted by hydrophobic residues like L104, F84 and L171. In the case of L108G, the substrate interacts more towards L104 than F84, which is the opposite in the rest of the variants **(Fig. S19d)**. The difference in the conformations of the substrate and the residues around it in L108G is detrimental for both the current step and the succeeding steps of the reaction. In the hyperactive variants, the phenyl group of the substrate shows more balanced hydrophobic interactions with residues around it. Moreover, closer proximity to the MIO cofactor gives the hyperactive variants a better chance in proceeding to the next step of the reaction when compared to the AvPAL* and L108G mutant.

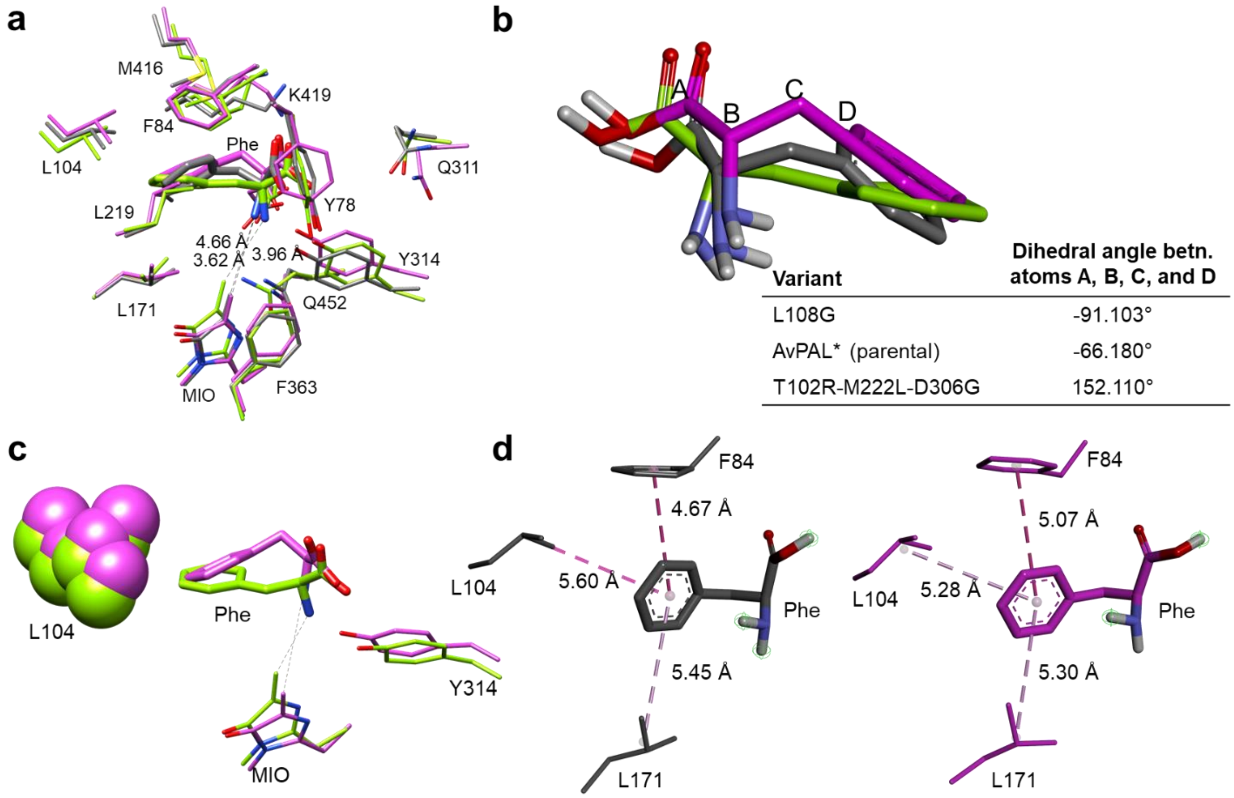

| Variant | Dihedral angle betn. atoms A, B, C, and D |
|---|---|
| L108G | -91.103° |
| AvPAL* (parental) | -66.180° |
| T102R-M222L-D306G | 152.110° |

**Figure S19. Observations of QM/MM studies. a)** Conformational changes of the active site residues and substrate for parental (grey), T102R-M222L-D306G (light green), and L108G (pink) are shown. The active site architecture was reconfigured after the mutation which yields a different conformation of substrate. **b)** The changes in dihedral angle between atoms C(A), CA(B), CB(C), and CG(D) of substrate were observed in parental, parental (grey), T102R-M222L-D306G (light green), and L108G (pink). **c)** L104, shown as space filling models, in T102R-M222L-D306G and L108G mutants was found to be occupying space in the pocket resulting the formation of non-catalytic conformation of Phe in L108G. **d)** The influence of residue L104 on Phe in wild and L108G mutant where, Phe is closer to L104 than F84 in L108G. The same is not seen in parental AvPAL* or the hyperactive variants. Non-polar hydrogens and hydrogen atoms on the active site residues are not displayed for ease of visualization.

**QM/MM metadynamics reveal different free energy paths to attain first Intermediate State (IS1).**

To understand the barrier-crossing events shown in the quantum mechanical/molecular mechanical (QM/MM) in depth, we combined hybrid QM/MM techniques combined with metadynamics[2], which enhanced the sampling of certain coordinates relevant to the reaction (termed collective variables, CVs). This way we could observe how the system accelerates across the reaction barriers by itself and escape from local minima (**Fig. S20**).

Also, the differences between AvPAL* variants can be revealed from the path taken by the reaction coordinates to attain the IS1; the main objective was to simulate the formation of an intermediate structure 1 (IS1) as described before in the reaction mechanism (**Fig. S16**). Two CVs important for the reaction were defined as follows; CV1: distance between the hydrogen of the amino nitrogen of Phe and hydroxyl oxygen of Y314, CV2: distance between the amino nitrogen of Phe and the methylidene carbon of MIO (**Table S5**). We further characterized the TS and FS (IS1) structures by transition path sampling simulations[3,4] and this was plotted over the FES (**Fig. S20**). In AvPAL* and the mutants, the attack conformation starts at 3.25 Å and moves as close as ~2.60 Å to form the first transition state (TS1). IS1 was observed in all the simulations but the minima differed, AvPAL* (CV1 = 3.0 Å, CV2 = 4.50 Å), L108G (CV1 = 3.0 Å, CV2 = 4.58 Å), M222L (CV1 = 2.80 Å, CV2 = 4.50 Å), G218S (CV1 = 3.0 Å, CV2 = 4.20 Å), T102E M222L (CV1=3.0 Å, CV2 = 4.60 Å), and T102R M222L D306G (CV1 = 3.0 Å, CV2 = 3.60 Å). The main difference was observed when the substrate approached MIO. For AvPAL* and L108G the barriers were prominent as the substrate approached 4.5 Å to MIO. In the other mutants, we could clearly see that there was more room for the substrate to advance closer to MIO, as this is required for next step of the reaction (**Fig. S20**). Moreover, the double and triple mutants showed minima as close as 3.5 Å.

These simulations demonstrate that the first step of the reaction is a proton abstraction occurs by tyrosine catalysed mechanism and this proton abstraction mechanism also is in line with experimental studies where the mutations M222L, G218S, T102E-M222L, T102R-M222L-D306G enhances this abstraction thereby making them hyperactive. These mutants follow a minimum energy path advancing closer to MIO that would facilitate the next step of the reaction

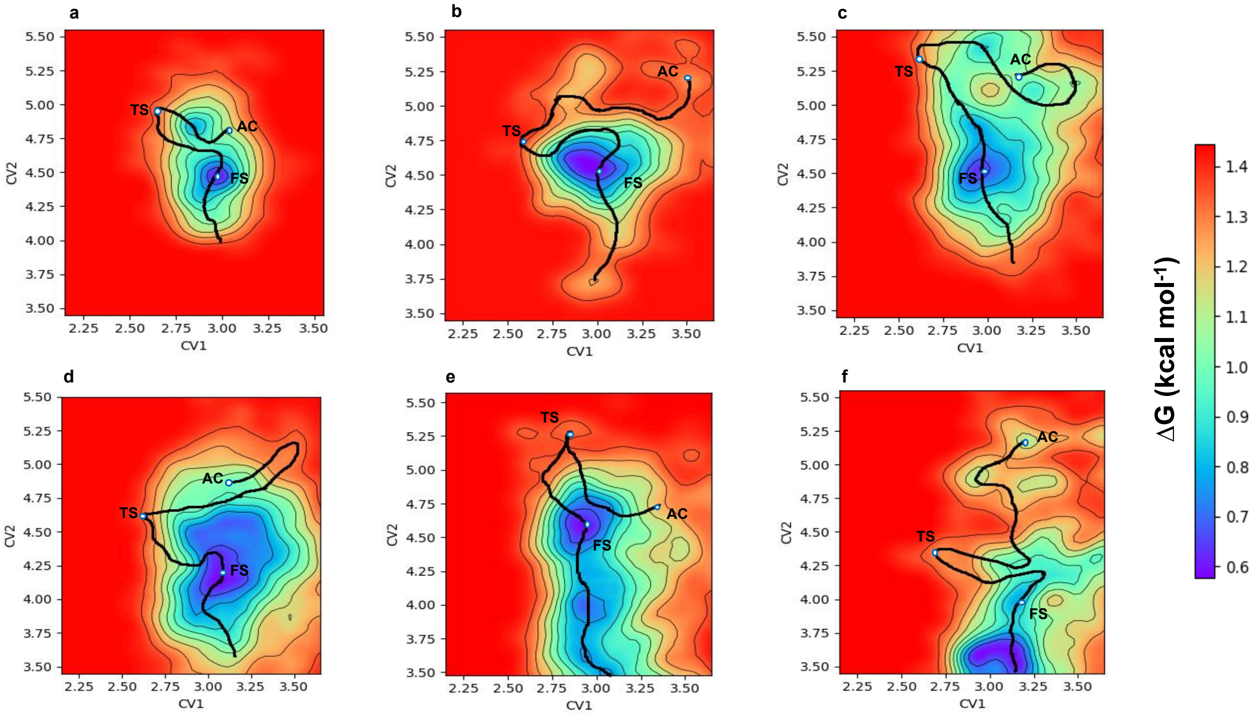

**Figure S20. Free energy surface (FES) from QM/MM metadynamics studies. a)** AvPAL*, **b)** L108G, **c)** M222L, **d)** G218S, **e)** T102E M222L, **f)** T102R M222L D306G, respectively. Trajectory obtained from the transition path sampling is plotted over the FES of all simulations. The attack conformation (AC), transition state (TS), and final state (FS) are labelled and shown as white circles. Two collective variables were employed for the simulations, CV1 and CV2, as described in the text. The free energy path for the mutations were more feasible for the next step of the reaction wherein the amino group of the substrate forms a covalent bond with the cofactor MIO as shown in the reaction mechanism (**Fig. S10**).

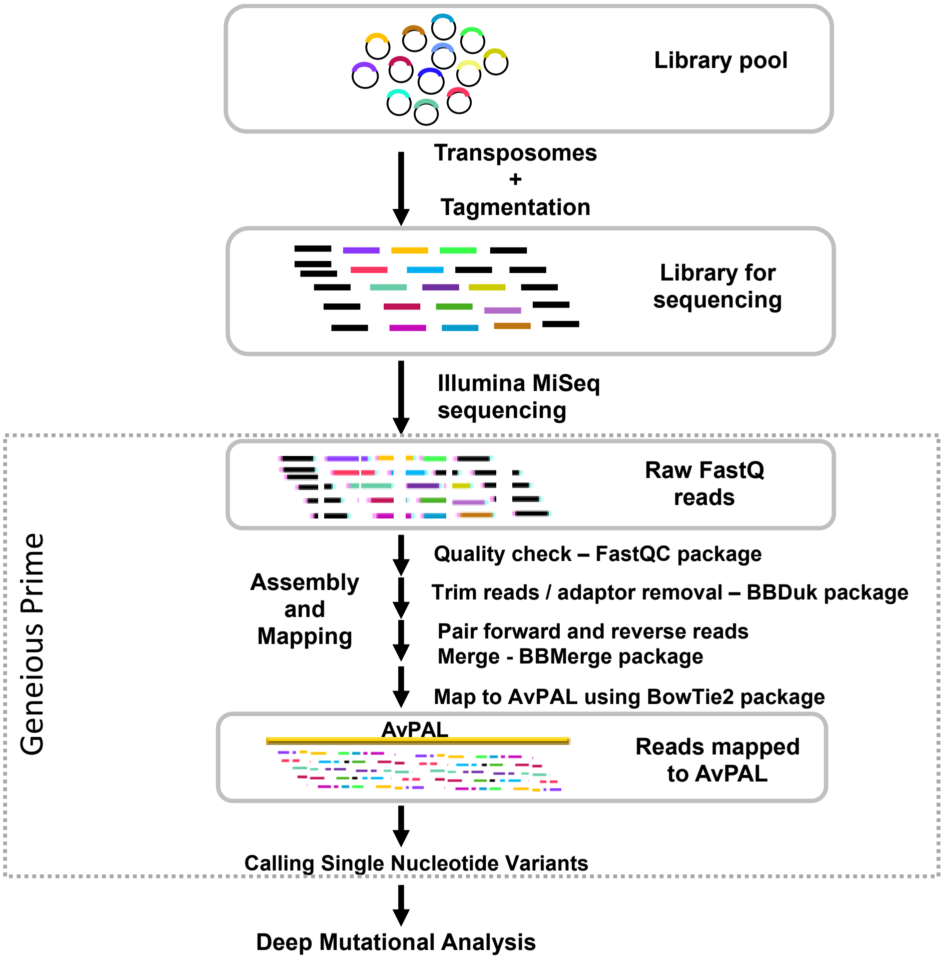

**Figure S21. Bioinformatic workflow.** Summary of workflow used for analysis of deep sequencing data of the libraries.

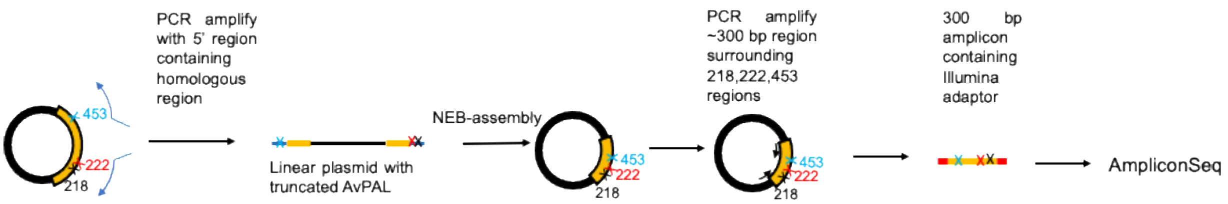

**Figure S22. Strategy for linking distal sites for amplicon sequencing.** The region immediate downstream to M222 in red, and immediate upstream to N453 in blue, was amplified with primers having homologous region. The amplicon flanked by homologous region was sealed using NEB-HiFi assembler. The circular plasmid was then used as a template to amplify ~300 bp region spanning G218-N453. The segment was amplified using primers with illumine sequencing overhangs. The amplicon was sequenced using AmpliconEZ Seq Illumina platform at Genewiz.

### SUPPLEMENTARY TABLES.

**Table S1.** Amino acid variation observed at L108 in AvPAL* homologs.

| AA substitution | Accession number | Organism | Function |
|---|---|---|---|
| Q | WP_182813033 | *Bacillus sp.* | Aromatic ammonia lyase |
| | WP_196473224 | *Bacillus sp.* | Aromatic ammonia lyase |
| | WP_074603200 | *Bacillus cereus* | Aromatic ammonia lyase |
| | XP_033537065 | *Eremomyces bilateralis* | Phenylalanine ammonia lyase |
| | XP_007812342 | *Metarhizium acridum* | Phenylalanine ammonia lyase |
| A | XP_033408294 | *Arthroderma uncinatum* | Phenylalanine ammonia lyase |
| K | WP_086933918 | *Agarilytica rhodophyticola* | Aromatic ammonia lyase |
| | WP_123638101 | *Marinimicrobium koreense* | Aromatic ammonia lyase |
| M | WP_046849712 | *Nitrosomonas communis* | Aromatic ammonia lyase |
| | WP_074666386 | *Nitrosomonas communis* | Aromatic ammonia lyase |
| | WP_132692682 | *Rubrobacter taiwanensis* | Aromatic ammonia lyase |
| | WP_105500951 | *Candidatus Sulfopaludibacter* | Aromatic ammonia lyase |
| T | XP_033437179 | *Daldinia childiae* | Phenylalanine ammonia lyase |
| | XP_002147032 | *Talaromyces marneffei* | Phenylalanine ammonia lyase |

**Table S2**: Frequency of different mutation combination occurring at G218-M222-N453 in Ep-PCR enriched library.

| | [G218S/A]-[M222L]-[N453S] (% frequency) | | | | | | | | | | | |
|---|---|---|---|---|---|---|---|---|---|---|---|---|
| | Wild type | Single mutations | | | | Double mutations | | | | | Triple mutations | |
| Passage | GMN* | SMN | AMN | GLN | GMS | AMS | ALS | SMS | SLS | GLS | ALN | SLN |
| #1 | 93.1 | 1.6 | 0.9 | 1.3 | 3.0 | 0.0 | 0.0 | 0.1 | 0.0 | 0.1 | 0.0 | 0.0 |
| #2 | 89.6 | 2.3 | 1.0 | 1.8 | 5.0 | 0.1 | 0.0 | 0.1 | 0.0 | 0.1 | 0.0 | 0.0 |
| #3 | 81.6 | 5.4 | 2.5 | 2.7 | 6.9 | 0.2 | 0.0 | 0.4 | 0.0 | 0.2 | 0.0 | 0.0 |

* mutations given as triplets at positions 218-222-453. WT is G218-M222-N453 and thus, GMN.

**Table S3.** Kinetic constants of AvPAL* variants combined with N453S.

| **PAL-variant** | **Model** | **$V_{max}$ (μmole·min$^{-1}$·mg$^{-1}$)** | **$K_M$ (μM)** | **$K_i$ (μM)** | **$k_{cat}$ (s$^{-1}$)** | **$k_{cat}/K_M$ (s$^{-1}$·μM$^{-1}$)** |
|---|---|---|---|---|---|---|
| AvPAL* | MM | 0.93 ± 0.01 | 137 ± 0 | - | 0.97 | 0.007 |
| N453S | MM | 0.85 ± 0.01 | 175 ± 15 | - | 0.89 | 0.005 |
| M222L | MM | 1.85 ± 0.02 | 145 ± 7 | - | 1.9 | 0.013 |
| M222L-N453S | SI | 1.01 ± 0.02 | 113 ± 8 | 167 ± 42 | 1.0 | 0.009 |
| L4P-G218S | SI | 1.86 ± 0.02 | 199 ± 9 | 208 ± 39 | 1.9 | 0.010 |
| L4P-G218S-N453S | - | - | - | - | - | - |
| T102E-M222L | MM | 2.33 ± 0.02 | 144 ± 6 | - | 2.4 | 0.017 |
| T102E-M222L-N453S | SI | 1.01 ± 0.02 | 92 ± 10 | 165 ± 57 | 1.1 | 0.011 |
| T102R-M222L-D306G | SI | 2.16 ± 0.02 | 96 ± 4 | 169 ± 21 | 2.3 | 0.023 |
| T102R-M222L-D306G-N453S | - | - | - | - | - | - |

MM – Michaelis-Menten, SI – Substrate inhibition (**Eqn S1, S2**)

**Table S4.** Growth rate of AvPAL* and N453S variants.

| **Variant** | **μmax (h$^{-1}$)** |
|---|---|
| AvPAL* | 0.14 |
| N453S | 0.175 |
| T102E-M222L | 0.183 |
| T102E-M222L-N453S | 0.20 |

**Table S6**. Michaelis complex (near attack conformation), derived from MD simulations.

| **Variant** | **Y314(O)−Phe(N) in Å** |
|---|---|
| AvPAL* | 2.497 |
| M222L | 2.494 |
| G218S | 2.486 |
| N453S | 2.485 |
| L108G | 2.510 |
| T102E-M222L | 2.473 |
| T102R-M222L-D306G | 2.472 |

**Table S7.** Attack conformation derived from QM/MM simulation

| Variant | Y314(O)−Phe($NH_3$, closest H) in Å |
|---|---|
| AvPAL* | 1.49 |
| G218S | 1.30 |
| M222L | 1.41 |
| N453S | 1.57 |
| L108G | 1.47 |
| T102E-M222L | 1.51 |
| T102R-M222L-D306G | 1.46 |

**Table S8.** Codons used for $^{7}C_{7}$ library.

| Position | Codon | Amino acids | # Codons | # AA |
|---|---|---|---|---|
| 102 | RNG | A, E, G, K, M, R, T, V | 8 | 8 |
| 218 | NNS | A, C, D, E, F, G, H, I, K, L, M, N, P, Q, R, S, T, V, W, Y, STOP | 32 | 21 |
| 222 | NNS | A, C, D, E, F, G, H, I, K, L, M, N, P, Q, R, S, T, V, W, Y, STOP | 32 | 21 |
| 268 | VNM | A, D, E, G, H, I, K, L, N, P, Q, R, S, T, V | 24 | 15 |
| 306 | GRC | D, G | 2 | 2 |
| 360 | VNR | A, E, G, I, K, L, M, P, Q, R, T, V | 24 | 12 |
| 453 | DDS | C, D, E, F, G, I, K, L, M, N, R, S, V, W, Y, STOP | 18 | 16 |

#### SUPPLEMENTARY EQUATIONS.

**Equation S1:**

Michaelis-Menten: $v = \frac{v_{max}[S]}{K_m+[S]}$

**Equation S2:**

Substrate inhibition: $v = \frac{v_{max}[S]}{K_m+[S]\left(1+\frac{[S]}{K_i}\right)}$

where,

$v$ = initial velocity,

$v_{max}$ = maximum initial velocity,

$K_m$ = Michaelis constant,

$K_i$ = inhibition constant for substrate

### SUPPLEMENTARY REFERENCES.

1 Jun, S.-Y. *et al.* Biochemical and structural analysis of substrate specificity of a phenylalanine ammonia-lyase. *Plant physiology* **176**, 1452-1468 (2018).

2 McGrath, M. J., Kuo, I.-F. W., Hayashi, S. & Takada, S. Adenosine triphosphate hydrolysis mechanism in kinesin studied by combined quantum-mechanical/molecular-mechanical metadynamics simulations. *Journal of the American Chemical Society* **135**, 8908-8919 (2013).

3 Dellago, C., Bolhuis, P. G. & Chandler, D. Efficient transition path sampling: Application to Lennard-Jones cluster rearrangements. *The Journal of chemical physics* **108**, 9236-9245 (1998).

4 Bolhuis, P. G., Chandler, D., Dellago, C. & Geissler, P. L. Transition path sampling: Throwing ropes over rough mountain passes, in the dark. *Annual review of physical chemistry* **53**, 291-318 (2002).
